# Genetic analysis of praziquantel response in schistosome parasites implicates a Transient Receptor Potential channel

**DOI:** 10.1101/2021.06.09.447779

**Authors:** Winka Le Clec’h, Frédéric D. Chevalier, Ana Carolina A. Mattos, Amanda Strickland, Robbie Diaz, Marina McDew-White, Claudia M. Rohr, Safari Kinung’hi, Fiona Allan, Bonnie L Webster, Joanne P Webster, Aidan M Emery, David Rollinson, Amadou Garba Djirmay, Khalid M Al Mashikhi, Salem Al Yafae, Mohamed A Idris, Hélène Moné, Gabriel Mouahid, Philip LoVerde, Jonathan S. Marchant, Timothy J.C. Anderson

## Abstract

Mass treatment with praziquantel (PZQ) monotherapy is the mainstay for schistosomiasis treatment. This drug shows imperfect cure rates in the field and parasites showing reduced PZQ response can be selected in the laboratory, but the extent of resistance in *Schistosoma mansoni* populations is unknown. We examined the genetic basis of variation in PZQ response in a *S. mansoni* population (SmLE-PZQ-R) selected with PZQ in the laboratory: 35% of these worms survive high dose (73 µg/mL) PZQ treatment. We used genome wide association to map loci underlying PZQ response. The major chr. 3 peak contains a transient receptor potential (*Sm.TRPM_PZQ_*) channel (Smp_246790), activated by nanomolar concentrations of PZQ. PZQ response shows recessive inheritance and marker-assisted selection of parasites at a single *Sm.TRPM_PZQ_* SNP enriched populations of PZQ-resistant (PZQ-ER) and sensitive (PZQ-ES) parasites showing >377 fold difference in PZQ response. The PZQ-ER parasites survived treatment in rodents better than PZQ-ES. Resistant parasites show 2.25-fold lower expression of *Sm.TRPM_PZQ_* than sensitive parasites. Specific chemical blockers of *Sm.TRPM_PZQ_* enhanced PZQ resistance, while *Sm.TRPM_PZQ_* activators increased sensitivity. A single SNP in *Sm.TRPM_PZQ_* differentiated PZQ-ER and PZQ-ES lines, but mutagenesis showed this was not involved in PZQ-response, suggesting linked regulatory changes. We surveyed *Sm.TRPM_PZQ_* sequence variation in 259 parasites from the New and Old World revealing one nonsense mutation that results in a truncated protein with no PZQ-binding site. Our results demonstrate that *Sm.TRPM_PZQ_* underlies variation in PZQ response in *S. mansoni* and provides an approach for monitoring emerging PZQ-resistance alleles in schistosome elimination programs.

**One Sentence Summary:** A transient receptor potential channel determines variation in praziquantel-response in *Schistosoma mansoni*.

## INTRODUCTION

Praziquantel (PZQ) is the drug of choice for treating schistosomiasis, a snail vectored parasitic disease, caused by flatworms in the genus *Schistosoma*. Schistosomiasis is widespread: three main parasite species infect over 140 million people in Africa, the Middle-East, South America and Asia (*1, 2*), resulting in widespread morbidity – a global burden of 1.9 million disability adjusted life years (*3*) – and mortality estimates ranging from 20,000 to 280,000 people annually (*4, 5*). Pathology results from eggs that lodge in the liver and intestine (*S. mansoni* and *S. japonicum*) or in the urogenital system (*S. haematobium*) stimulating granuloma formation. This results in a spectrum of pathology including portal hypertension, hepatosplenic disease, bladder cancer, genital schistosomiasis and infertility. *S. mansoni* infection alone results in a conservative estimate of 8.5 million cases of hepatosplenomegaly in sub-Saharan Africa (*6*). Mass drug administration programs currently distribute an estimated 250 million doses of PZQ per year aimed in the short term at reducing schistosome associated morbidity and mortality, and in the longer term at eliminating schistosomiasis transmission (*7, 8*). PZQ is also widely used for treatment of other flatworm parasites of both humans and livestock including tapeworms.

PZQ treatment of adult worms results in rapid Ca^2+^ influx into cells, muscle contraction and tegument damage (*9–12*). Both the mechanism of action and the mechanism of resistance to PZQ have been the focus for much speculation and research (*13, 14*). Several proteins like voltage-gated calcium channels (*15–17*) or ABC transporters (*18, 19*) have been suspected to play a role in PZQ resistance. However, this topic has been stimulated by the recent finding that a transient receptor channel (*Sm.TRPM_PZQ_*) is activated by nanomolar quantities of PZQ (*20, 21*).

Mass drug treatment with PZQ has enormous health benefits and has been extremely effective in reducing parasite burdens and transmission (*8*), but imposes strong selection for resistance on treated schistosome populations. Emergence of PZQ resistance is a major concern, because it could derail current progress towards the WHO goal of eliminating schistosomiasis as a public health problem by 2025 (*8*). Several lines of evidence from both the field and the laboratory suggest that PZQ response varies in schistosome populations (*22–27*). PZQ resistance is readily selected in the laboratory through treatment of infected rodents or infected intermediate snail hosts (*28*). This typically results in a modest change (3-5 fold) in PZQ response in parasite populations (*28*), although the PZQ resistance status of individual worms comprising these populations is unknown. PZQ treatment typically results in ∼30% of patients who remain egg positive following treatment (*29*). PZQ kills adult worms, but not immature parasites (*30, 31*), so both newly emerging adult parasites and drug resistance may contribute to treatment failure. There have been several reports of patients who remained egg positive across multiple PZQ treatment cycles (*32, 33*): schistosome infections established in mice from infective larvae from these patients showed elevated resistance to PZQ (*24, 34*). In Kenya and Uganda, infected communities where prevalence and disease burden are not reduced by repeated treatment have been identified. The causes of these “hotspots” (*35, 36*) is currently unknown, but PZQ-resistant schistosomes are one explanation. In a large longitudinal study of individual school age children, egg reduction ratios (ERR) were high in naïve populations treated with PZQ, but showed a significant decline after multiple rounds of treatment (*37*), consistent with selection of tolerance or resistance to PZQ. Identification of molecular markers for direct screening of levels of PZQ resistance alleles would be extremely valuable for parasite control programs, because changes in schistosome ERR have both genetic and non-genetic explanations and are laborious to measure.

The availability of good genome sequence and near complete genome assembly (*38*) for *S. mansoni* make unbiased genome wide approaches feasible for schistosome research (*39*). Our central goal is to determine the genetic basis of variation in PZQ response, using genome wide association approaches. We exploit the PZQ-resistant parasites generated by laboratory selection (*26*) to determine the genetic basis of PZQ, identifying a transient receptor channel as the cause of variation in PZQ response. The Transient Receptor Potential Melastatin (TRPM) ion channel identified is activated by PZQ (*20, 21*) and has been designated *Sm.TRPM_PZQ_*. Together, our genetic analysis and the independent pharmacological analysis by Park *et al.* (*21*) identify the target channel for PZQ and provide a framework for monitoring PZQ resistance evolution in schistosomiasis control programs.

## RESULTS

### PZQ resistant parasites are present in laboratory schistosome populations

Male and female schistosome parasites pair in the blood vessels and reproduction is obligately sexual, so schistosomes are maintained in the laboratory as outbred populations. Hence, individual parasites within laboratory populations may vary in PZQ response. We measured PZQ response in the SmLE-PZQ-R population, which was previously generated by PZQ treatment of infected snails (*26*). This revealed a 14-fold difference in IC_50_ between the SmLE progenitor parasite population (IC_50_=0.86 ± 0.14 µg/mL) and SmLE-PZQ-R (IC_50_=12.75 ± 4.49 µg/mL, χ^2^ test, p = 0.001) derived by PZQ selection (Fig. 1). This is higher than the 3-5 fold differences observed between PZQ-selected and unselected parasite populations in previous studies (*40*). Interestingly, the dose response curve for the SmLE-PZQ-R population plateaus at 65% mortality: the remaining 35% of parasites recovered even at high dose of PZQ (72.9 µg/mL). These results suggest that the SmLE-PZQ-R parasite population is a mixed population that contains both PZQ-sensitive and PZQ-resistant individual worms (Fig. 1).

**Fig. 1.**
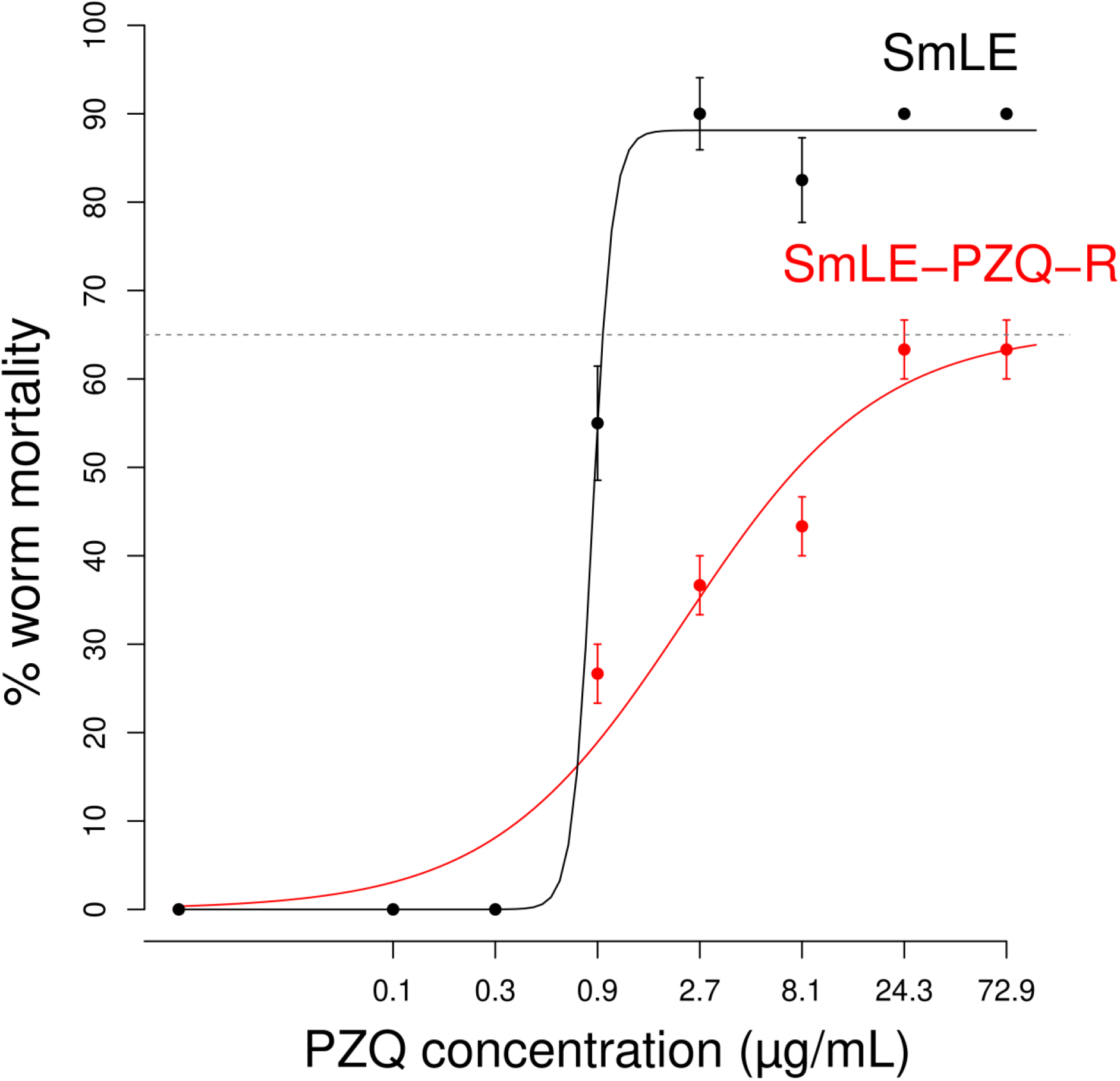
Dose response curves for SmLE (PZQ-S) and the derived SmLE-PZQ-R (PZQ-R) populations. PZQ dose response curves show a ∼14-fold difference in response between SmLE (ancestral population) and SmLE-PZQ-R (PZQ selected population) (χ^2^ test = 10.387, p = 0.001). The PZQ-selected laboratory schistosome population (SmLE-PZQ-R) is polymorphic for drug response. 35% of SmLE-PZQ-R are not killed by treatment with high dose of PZQ, suggesting that this population is polymorphic (*N*=240 worms/populations).

### Association mapping of PZQ resistant genes identifies a TRPM channel

We conducted a genome wide association study (GWAS) to determine the genetic basis of PZQ resistance (PZQ-R). GWAS has been widely used for mapping drug resistance in parasitic protozoa (*41*) and the model nematode *Caenorhabditis elegans* (*42*), but has not previously been applied to parasitic helminths, because of the difficulty of accurately measuring drug response in individual parasites. When worms are treated with PZQ, there is a massive influx of Ca^2+^ into cells and parasites contract (*17, 43*), but some worms recover and resume respiration and movement 24-48h after drug removal. We assayed parasite recovery following high dose PZQ treatment (24 µg/mL) of individual male worms maintained in 96-well plates by measuring L-lactate production (*44*), a surrogate measure of respiration, 48h after PZQ treatment removal (Fig. S1). These assays allow efficient measurement of recovery in individual PZQ-treated worms.

We conducted replicate experiments (A: n = 590; B: n = 691) to measure PZQ response in individual parasites maintained in 96-well plates. The distributions of L-lactate production in the two experiments were broad (A: 0-126.56 nmol/h, mean = 42.95 nmol/h; B: 0-118.61 nmol/h, mean = 37.79 nmol/h): we identified worms from the top and bottom quintile for L-lactate production (Fig. 2A) which were then bulk sequenced to high read depth (average read depth -A: 39.97; B: 36.83). Two genome regions (chr. 2 and chr. 3) showed strong differentiation in allele frequencies between parasite populations showing high and low L-lactate production phenotypes (Fig. 2B). The highest peak (p = 1.41 × 10^-22^) on chr. 3 spanned 4 Mb (22,805-4,014,031 bp) and contained 91 genes, of which 85 are expressed in adult worms (Fig. 2C). This genome region contains several potential candidate loci including three partial ABC transporters and a voltage-gated calcium channel subunit (Table S1). One gene close to the highest association peak is of particular interest: Smp_246790 is a transient receptor potential channel in the M family (TRPM). This same channel was recently shown to be activated by PZQ following exposure to nanomole quantities of drug resulting in massive Ca^2+^ influx into HEK293 cells transiently expressing this protein (*20, 21*). This gene, designated *Sm.TRPM_PZQ_*, is therefore a strong candidate to explain variation in PZQ response within parasite populations. Two other features of the data are of interest. First, the SNP (position 1029621 T>C) marking the highest association peak (at 1,030 Mb) is found in a transcription factor (Smp_345310, SOX13 homology) from a family known to regulate splicing variants (*45*). Second, there is a ∼150 kb deletion (1220683-1220861 bp) 6.5 kb from *Sm.TRPM_PZQ_* and another 150 kb deletion (1,200,000-1,350,000 bp) 170 kb from the transcription factor. This was enriched in high L-lactate groups in both replicates and is in linkage disequilibrium with the enriched SNP in *Sm.TRPM_PZQ_*.

**Fig. 2.**
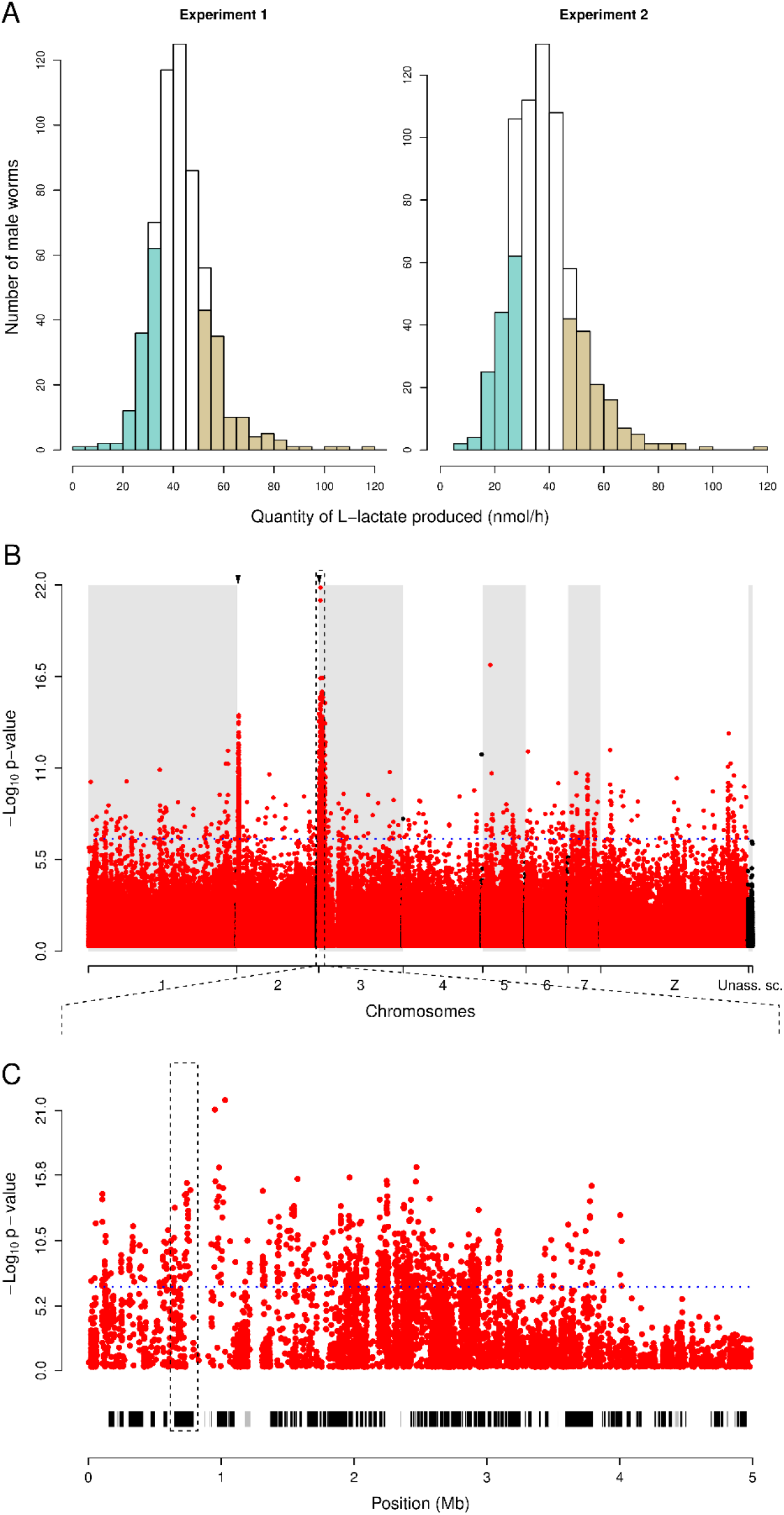
Genome-wide association mapping of PZQ response. **(A)** We measured recovery of individual adult male worms following expose to 24 µg/mL PZQ by measuring L-lactate production. The distribution from both experimental replicates is shown (A: *N*=590; B: *N*=691). Worms in the bottom (teal) and top (gold) quintile were each pooled, and genome sequenced to high read depth. **(B)** The Manhattan plot identifies genome regions that differ in allele frequency between high and low L-lactate worm pools. Blue dotted line refers to the Bonferroni significance threshold; red dots represent association of individual SNPs; orange arrows mark the position of prominent QTLs. **(C)** The chr. 3 QTL identified spans 4 Mb and 91 genes. Boxes under the Manhattan plot show gene locations (black = expressed, grey = unexpressed). Position of the Sm.TRPM_PZQ_ is marked with dashed lines.

The chr. 2 peak (p < 1.0 × 10^-15^) spans 1.166 Mb (291,191-1,457,462 bp) and contains 24 genes (21 expressed in adult worms) (Table S1). This genome region does not contain obvious candidate genes that might explain variation in PZQ response.

### PZQ resistance shows recessive inheritance

To confirm these associations and determine whether the loci underlying PZQ response are inherited in a dominant, co-dominant or recessive manner, we compared genotype and PZQ-response phenotype in individual worms. We compared the L-lactate production phenotypes of individual worms maintained in 96-well plates 48 hours after exposure to 24 µg/mL PZQ with their genotypes at SNPs at the peaks of the chr. 2 and chr. 3 QTLs. We also examined copy number of one of the 150 kb deletion observed on chr. 3 using qPCR. We observed significant differences in L-lactate production among genotypes (Fig. 3). Both the SNP assayed in *Sm.TRPM_PZQ_* and the copy number variant revealed that the causative gene in the chr. 3 QTL showed recessive inheritance (Fig 3B-C). Homozygous parasites carrying two copies of the *Sm.TRPM_PZQ_*-741987C allele (or two copies of the deletion) recovered from PZQ treatment, while the heterozygotes and other homozygotes failed to recover from treatment (Fig. 3B-C). For the chr. 2 QTL, we did not see a significant association between parasite genotype and PZQ-R phenotype nor with L-lactate production before PZQ treatment (Fig. 3A): this locus was not investigated further.

**Fig. 3.**
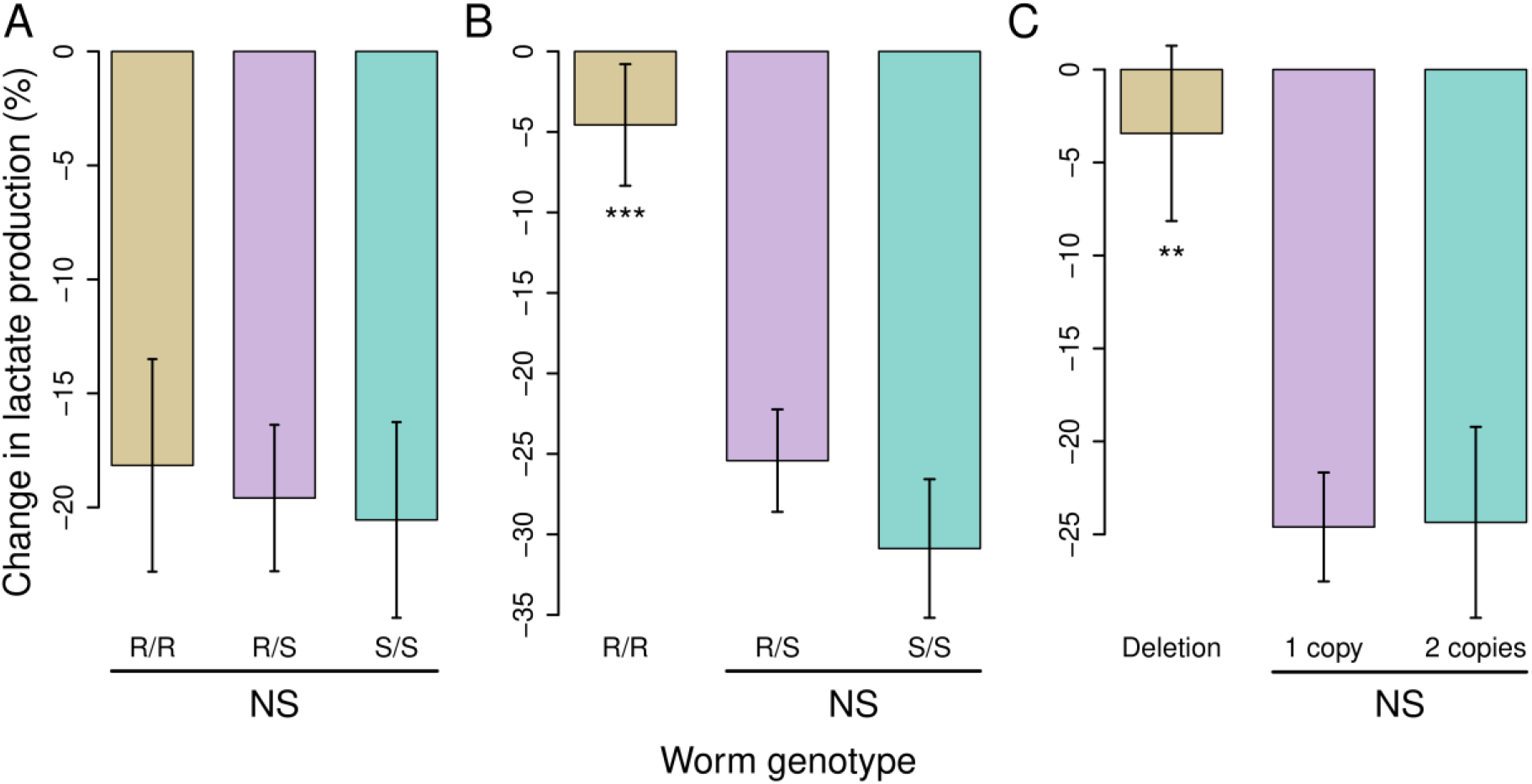
Inheritance of PZQ response in LE-PZQ population. Bar charts show the change in L-lactate production after exposure to 24 µg/mL PZQ in worms from different genotypic classes for QTL regions on chr. 2 and 3. **(A)** chr. 2 QTL (Kruskal-Wallis KW test χ^2^ = 0.019, p = 0.99), **(B)** *Sm.TRPM_PZQ_*-741987C (KW test χ^2^ = 24.481, p = 2.93×10^-6^), **(C)** 100kb deletion (KW test χ^2^ = 15.708, p = 0.0004). We see minimal change in L-lactate production following PZQ exposure in homozygotes for the SNP enriched in PZQ treated parasites, indicating that this trait is recessive. Parasites carrying two copies of the 100 kb deletion are also strongly associated with resistance, demonstrating that this deletion is in LD (*N* = 120 worms; NS: No significant difference between groups; \**p* < 0.05; ** *p* ≤ 0.01; *** *p* ≤ 0.001).

### Marker-assisted purification of resistant and sensitive parasites

As the chr. 3 QTL containing *Sm.TRPM_PZQ_* exhibits the strongest association with PZQ response and shows recessive inheritance, we were able to use single generation marker assisted selection approach to enrich parasites for alleles conferring PZQ resistance (PZQ-R) and PZQ sensitivity (PZQ-S) from the mixed genotype SmLE-PZQ-R parasite population (Fig. 4A). We genotyped clonal cercariae larvae emerging from snails previously exposed to single miracidia to identify parasites homozygous for the recessive PZQ-R allele from those homozygous for the PZQ-S allele. Parasites isolated from multiple snails falling into these two alternative genotypes were then used to infect hamsters. The enriched PZQ-resistant and sensitive parasites were designated SmLE-PZQ-ER and SmLE-PZQ-ES. Sequencing of adult parasites recovered from these two populations revealed that they were fixed for alternative alleles at the *Sm.TRPM_PZQ_*-741987C SNP genotyped, but showed similar allele frequencies across the rest of the genome (Fig. S2). As expected, these sequences also revealed that the 100 kb deletion was close to fixation in the SmLE-PZQ-ER population (Fig. S3).

**Fig. 4.**
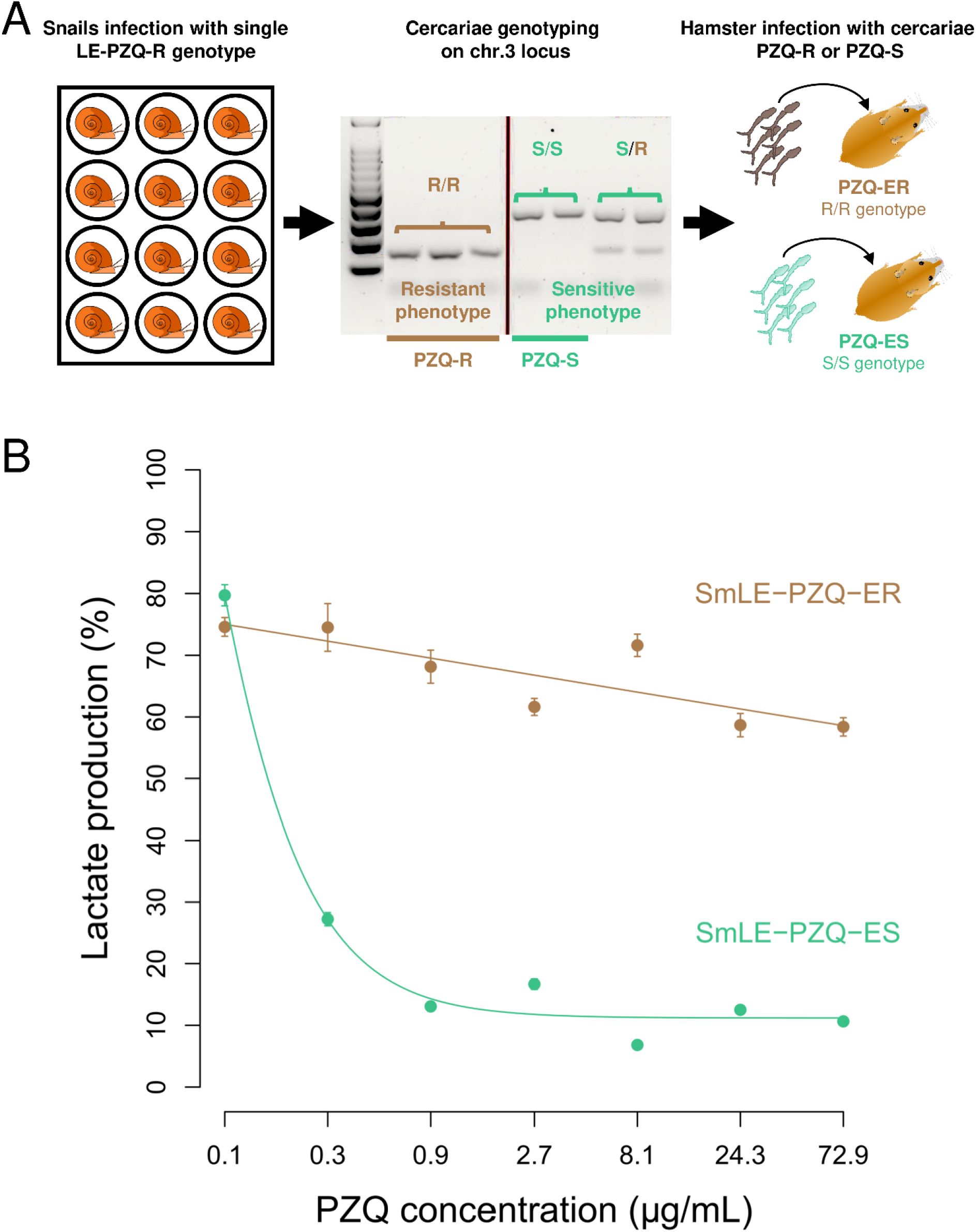
Single generation marker-assisted purification of SmLE-PZQ-ES and SmLE-PZQ-ER parasites. **(A)** Experimental strategy for identifying parasite larvae that are homozygous for *Sm.TRPM_PZQ_* alleles associated with PZQ-R or PZQ-S. We genotyped cercaria larvae emerging from snails infected with single parasite genotypes for a restriction site in the *Sm.TRPM_PZQ_* gene, and then infected two groups of hamsters with parasites homozygous for alternative alleles at this locus. **(B)** The two populations of parasites generated show dramatic differences in PZQ-response (*N* = 60 worms/population/treatment, χ^2^ test = 373.03, p < 2.2×10^-16^).

We conducted PZQ dose response curves on these enriched parasite populations. The SmLE-PZQ-ES population had an IC_50_ of 0.198 µg/mL (± 1 s. d.: 0.045), while the SmLE-PZQ-ER population did not reach 50% reduction even at the highest dose (72.9 µg/mL) so has an IC_50_ >72.9 µg/mL: the two purified populations differ by >377-fold in PZQ response (Fig. 4B). These results provide further demonstration that the original SmLE-PZQ-R parasite population was a mixture of PZQ-R and PZQ-S parasites. Separation of the component SmLE-PZQ-ES and SmLE-PZQ-ER parasites from these mixed populations allows rigorous characterization of the PZQ-R trait in parasite populations that are fixed for alternative alleles at *Sm.TRPM_PZQ_*, but contain comparable genomic backgrounds across the rest of the genome.

### *Sm.TRPM_PZQ_* gene and isoform expression is reduced in SmLE-PZQ-ER parasites

PZQ response varies between parasite stages and sexes, with strongest response in adult males. Adult females and juvenile worms are naturally less susceptible (*30*). We therefore examined gene expression in the purified SmLE-PZQ-ES and SmLE-PZQ-ER populations (males and females for both adult and juvenile worms) using RNA-seq (Fig. 5). Of the 85 genes expressed in adult worms under the chr. 3 QTL, only the *Sm.TRPM_PZQ_* showed a significant reduction in expression in the SmLE-PZQ-ER adult male worms relative to SmLE-PZQ-ES (2.25-fold, posterior probability = 1) (Fig. 5A-B). Comparable under expression of *Sm.TRPM_PZQ_* was also seen in female worms when compared to SmLE-PZQ-ES: expression of *Sm.TRPM_PZQ_* was 11.94-fold lower in female than in male worms, consistent with females being naturally resistant (*30*) (Fig. 5C). However, juvenile male and female worms showed elevated gene expression compared with adult worms (Fig. 5C). This is surprising because juveniles are naturally resistant to PZQ. *Sm.TRPM_PZQ_* has 41 exons and occurs as 7 isoforms containing between 3 and 36 exons. Strikingly, SmLE-PZQ-ES male worms showed a 4.02-fold higher expression of isoform 6 compared to SmLE-PZQ-ER males, and an 8-fold higher expression than naturally resistant juvenile worms from both populations (while SmLE-PZQ-ER showed only a 2-fold higher expression) (Fig. 5B and D and Fig. S4). This suggests that high expression of isoform 6 is associated with PZQ sensitivity. The 15 exons of isoform 6 produce an 836 amino acid protein that lacks the transmembrane domain but contains the TRP domain. The function of isoform 6 is unclear and we don’t know if this association with PZQ sensitivity is causal. We interrogated the 10x single cell expression data from adult worms (*46*) showing that *Sm.TRPM_PZQ_* is expressed mainly in neural tissue with some expression also in muscle (Fig. S5).

**Fig. 5.**
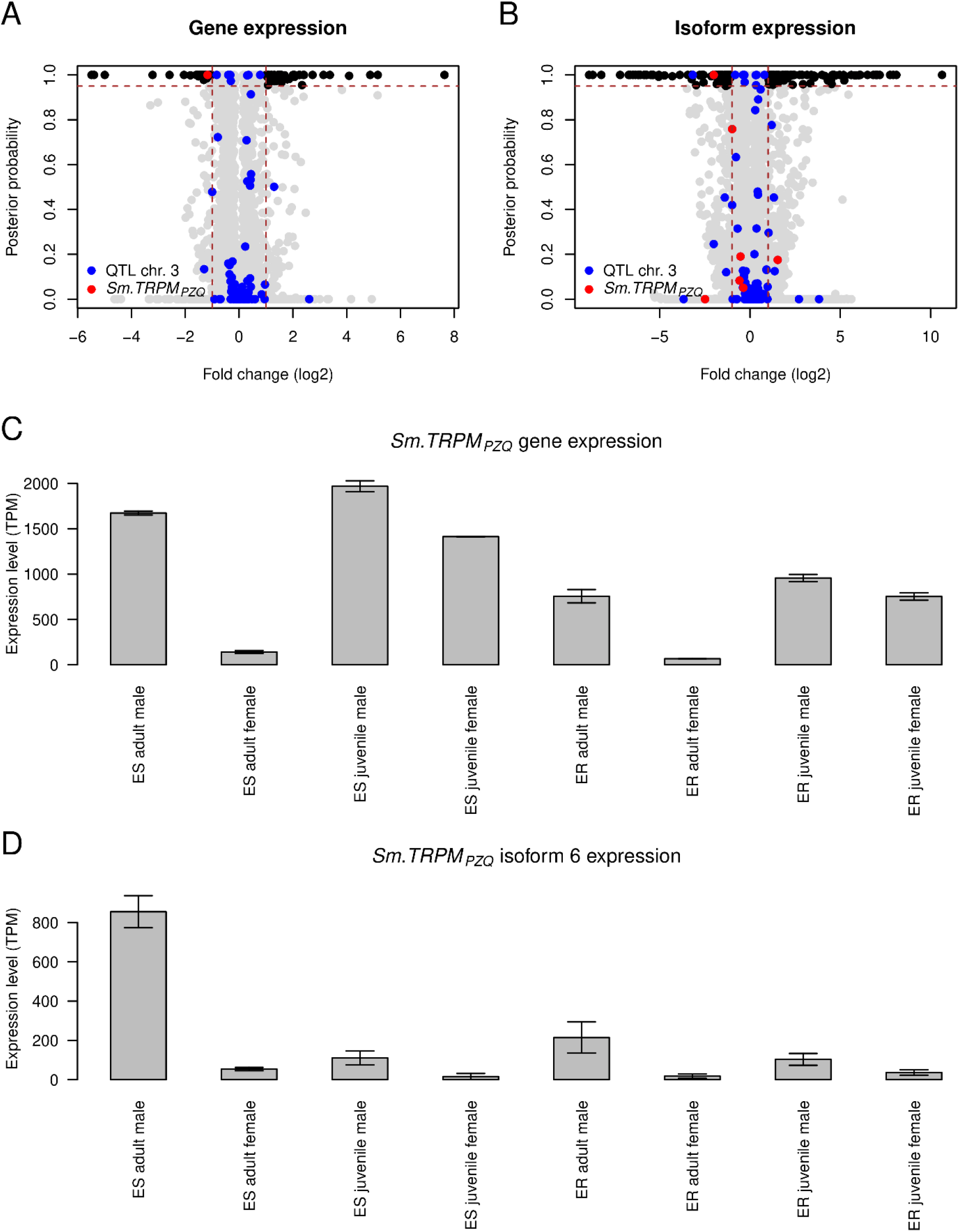
Gene expression differences between SmLE-PZQ-ES and SmLE-PZQ-ER parasites. Volcano plot showing the differential **(A)** gene expression and **(B)** isoform expression between adult male PZQ-ES and PZQ-ER (in black: genes or isoforms showing significant differential expression genome-wide, in blue: genes or isoforms located under the chr. 3 QTL, in red: *Sm.TRPM_PZQ_* gene or isoform). **(C)** *Sm.TRPM_PZQ_* gene expression and **(D)** *Sm.TRPM_PZQ_* isoform 6 expression level comparison between PZQ-ES and ER for the two sex (i.e. male and females) and different stages (i.e. adult and juvenile). High expression of *Sm.TRPM_PZQ_* isoform 6 is associated with PZQ sensitivity.

### Fitness of SmLE-PZQ-ER and SmLE-PZQ-ES parasite populations

Both laboratory selected and field isolated *S. mansoni* showing PZQ-R have been difficult to maintain in the laboratory (*47*): the PZQ-R trait has been rapidly lost consistent with strong selection against this trait. It has been suggested that PZQ-R carries a fitness cost that will slow spread of this trait in the field under PZQ pressure. Such fitness costs are a common, but not ubiquitous, feature of drug resistance in other pathogens (*48–50*). We measured several components of parasite fitness in SmLE-PZQ-ES and SmLE-PZQ-ER parasites during laboratory passage of purified parasite lines, but found no significant differences in infectivity to snails, snail survival, or infectivity to hamsters (Fig. S6). We did not see loss of PZQ-R in our lines after 12 generations because the key genome region is fixed. Cioli *et al.* (*40*) has also reported long term stability of PZQ-R parasite populations indicative that PZQ-R associated fitness costs maybe limited or absent.

### *In vivo* efficacy of PZQ against SmLE-PZQ-ES and SmLE-PZQ-ER parasites

To determine the relationship between *in vitro* PZQ-R measured in 96-well plates, and *in vivo* resistance, we treated mice infected with either SmLE-PZQ-ER or SmLE-PZQ-ES parasites populations with 120 mg/kg of PZQ. We observed no significant reduction in worm burden in SmLE-PZQ-ER parasites when comparing PZQ-treated and control (DMSO-treated) animals (Wilcoxon test, p = 0.393; Fig. S7A). In contrast, we recovered significantly lower numbers of worms from PZQ-treated versus untreated mice infected with the SmLE-PZQ-ES parasite population (Wilcoxon test, p = 0.008; Fig. S7A). The percent reduction observed was significantly different between the SmLE-PZQ-ES and SmLE-PZQ-ER parasites (Wilcoxon test, p = 0.0129; Fig. S7B). Interestingly, we observed a large reduction in numbers of female worms recovered from PZQ-treated SmLE-PZQ-ES parasites relative to untreated animals (Wilcoxon test, p = 0.008; Fig. S7D), while for male worms this did not reach significance (Wilcoxon test, p = 0.089, Fig. S7C). We saw no impact of PZQ-treatment for either female or male worms in mice infected with SmLE-PZQ-ER. These results show that *in vivo* PZQ response in treated mice differs between SmLE-PZQ-ES and SmLE-PZQ-ER parasites. These data also suggest that the extended paralysis of male SmLE-PZQ-ES worms under PZQ treatment may reduce their ability to maintain female worms *in copula*.

### Chemical blockers and activators of *Sm.TRPM_PZQ_* modulate PZQ-R

*Sm.TRPM_PZQ_* emerges as a strong candidate gene to explain variation in PZQ response, but validation is required. We were unsuccessful in knocking down expression of *Sm.TRPM_PZQ_* using either siRNA or dsRNA (Table S2), possibly because this gene is expressed mainly in neural tissue. We therefore used two chemical modulators of Sm.TRPM_PZQ_ activity – an Sm.TRPM_PZQ_ agonist (AG1) and Sm.TRPM_PZQ_ antagonist (ANT1). These were identified among ∼16,000 compounds by screening Ca^2+^ influx into HEK293 cells transiently expressing Sm.TRPM_PZQ_ (Chulkov *et al.*, in prep). Addition of the Sm.TRPM_PZQ_ blocker (ANT1) allowed SmLE-PZQ-ES worms to recover from PZQ treatment (Fig. 6A), while the Sm.TRPM_PZQ_ activator (AG1) rendered SmLE-PZQ-ER worms sensitive to PZQ treatment in a dose dependent manner (Fig. 6B). These results are consistent with a role for Sm.TRPM_PZQ_ in determining variation in PZQ response.

**Fig. 6.**
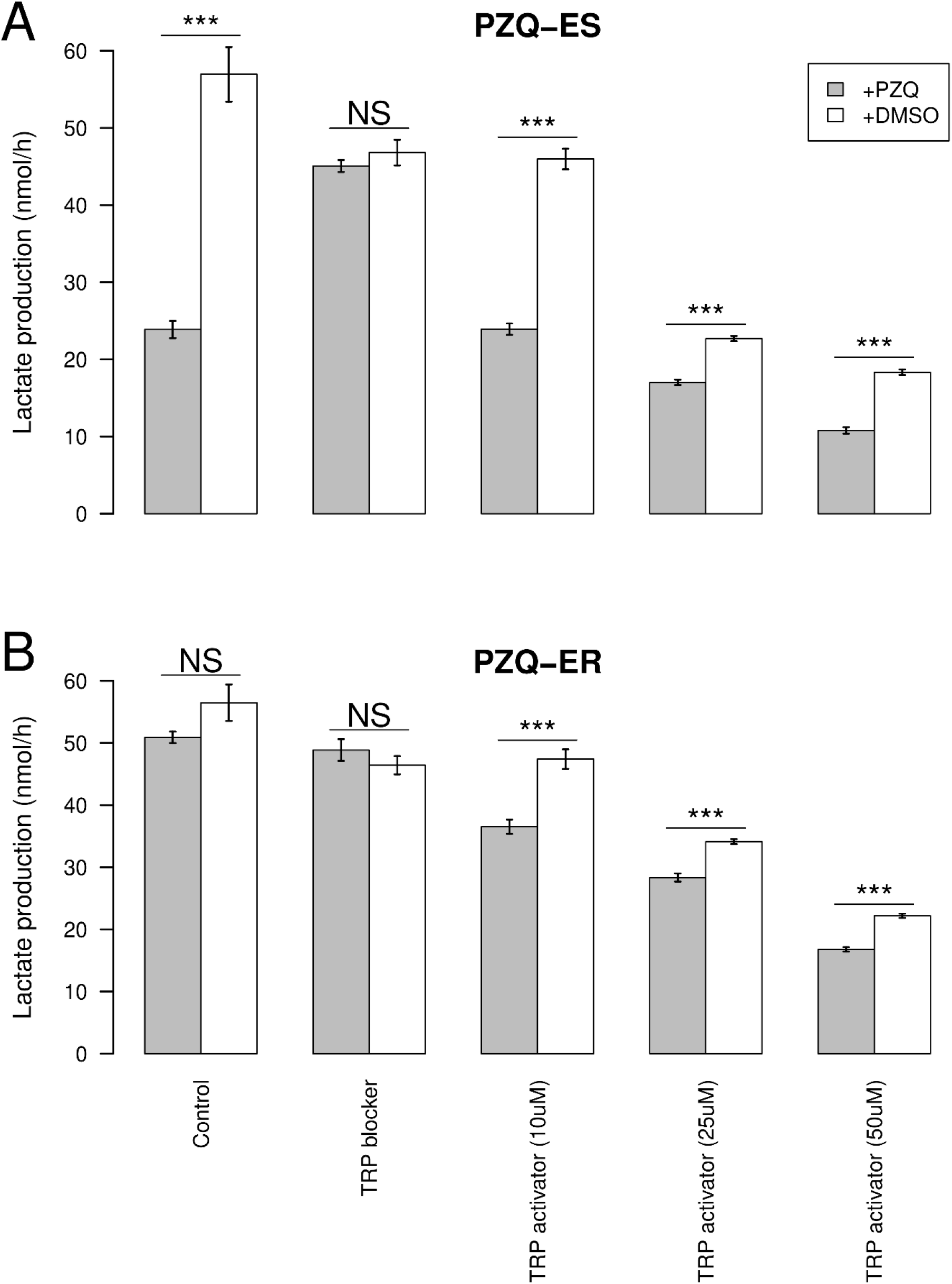
Impact of *Sm.TRPM_PZQ_* blockers and activators on PZQ response. **(A)** SmLE-PZQ-ES and **(B)** SmLE-PZQ-ER were exposed to either i) PZQ or DMSO alone (control group), ii) PZQ or DMSO combined with 50 µM Sm.TRPM_PZQ_ blocker (antagonist ANT1), iii) PZQ or DMSO combined with either 10 µM, 25 µM or 50 µM of Sm.TRPM_PZQ_ activator (agonist AG1). Parasite viability was assessed 3 days post-treatment, based on their L-lactate production. Addition of the Sm.TRPM_PZQ_ blocker allowed SmLE-PZQ-ES worms to recover from PZQ treatment (Welsh t-test, t = −0.94, p = 0.35), while the Sm.TRPM_PZQ_ activator (agonist AG1) rendered SmLE-PZQ-ER worms sensitive to PZQ treatment in a dose dependent manner (N = 20 worms/population/treatment; Welsh t-test, NS: No significant difference between groups; \**p* < 0.05; ** *p* ≤ 0.01; *** *p* ≤ 0.001.)

We found 5 non-synonymous SNPs and 5 insertions that showed significant differences in allele frequency in *Sm.TRPM_PZQ_* in the SmLE-PZQ-ES and SmLE-PZQ-ER parasite populations. One of these SNPs (*Sm.TRPM_PZQ_*-741903) and two insertions (*Sm.TRPM_PZQ_*-779355 and *Sm.TRPM_PZQ_*-779359) are fixed for alternative alleles in the two populations, with 7 others segregating at different frequencies in the two populations (Fig. S2). These SNPs are located outside the critical transmembrane domains so were not strong candidates to explain differences in PZQ-R. We expressed *Sm.TRPM_PZQ_* carrying some of these variants in HEK293 cells and examined their impact on Ca^2+^ influx to interrogate their role in explaining difference in PZQ response (Fig. S2). None of these SNPs affect Ca^2+^ influx: they are therefore unlikely to underlie PZQ response. We speculate that the difference in PZQ response is due to expression patterns and may be controlled by regulatory variants potentially associated with the adjacent 150 kb deletion or the SOX13 transcription factor.

### Sequence variation in *Sm.TRPM_PZQ_* from natural *S. mansoni* populations

Methods for evaluating frequencies of PZQ-resistance mutations in endemic regions would provide a valuable tool for monitoring mass treatment programs aimed at schistosome elimination. Both this paper and the accompanying paper (*21*) identify *Sm.TRPM_PZQ_* as being critical to PZQ-response, and Park *et al.* have determined critical residues that determine binding between PZQ and *Sm.TRPM_PZQ_* (*21*). We examined the mutations present in *Sm.TRPM_PZQ_* in natural schistosome populations using exome sequencing from 259 miracidia, cercariae or adult parasites from 3 African countries (Senegal, Niger, Tanzania), the Middle East (Oman) and South America (Brazil) (*51, 52*). We were able to sequence 36/41 exons of *Sm.TRPM_PZQ_* from 122/259 parasites on average (s.e. = 18.65) (Table S3). We found 1 putative PZQ-R SNP in our reads supported by a very high coverage (Fig. S8). This SNP (c.2708G>T on isoform 5, p.G903*) was found in a single Omani sample and resulted in a truncated protein predicted to result in loss-of-function, demonstrating that PZQ-R alleles are present in natural populations. However, this PZQ-R allele observed was rare and present in heterozygous state so would not impact PZQ response (Fig. S9).

## DISCUSSION

Our genetic approach to determining the genes underlying PZQ resistance – using GWAS and a simple lactate-based read out to determine parasite recovery following PZQ treatment in individual parasites – robustly identifies a TRPM channel (*Sm.TRPM_PZQ_*) as the cause of variation in PZQ response. We were further able to purify SmLE-PZQ-ER and SmLE-PZQ-ES parasites to examine drug response and gene expression and to use chemical blockers to directly implicate *Sm.TRPM_PZQ_*. Our results complement those of Park *et al.* (*21*) who used a pharmacological approach to determine that *Sm.TRPM_PZQ_* is the major target for PZQ, and identified the critical residues necessary for activation by PZQ. Together, these approaches demonstrate that TRPM is a key determinant of schistosome response to PZQ.

A striking feature of the results is the strength of the PZQ-R phenotype. While previous authors have described quite modest differences (3-5 fold) in PZQ-response among *S. mansoni* isolates (*40*), this study revealed at least 377-fold difference in IC_50_ between SmLE-PZQ-ER and SmLE-PZQ-ES parasites. These large differences were only evident after we used marker-assisted selection to divide a mixed genotype laboratory *S. mansoni* population into component SmLE-PZQ-ER and SmLE-PZQ-ES populations. The modest IC_50_ differences in previous studies observed are most likely because the parasite lines compared contained mixed populations of both SmLE-PZQ-ER and SmLE-PZQ-ES individuals. This highlights a critical feature of laboratory schistosome populations that is frequently ignored: these populations are genetically variable and contain segregating genetic and phenotypic variation for a wide variety of parasite traits. In this respect they differ from the clonal bacterial or protozoan parasite “strains” used for laboratory research. Importantly, we can use this segregating genetic variation for genetic mapping of biomedically important parasite traits such as PZQ resistance.

There is strong evidence that PZQ-R parasites occur in schistosome populations in the field, but the contribution of PZQ-R to treatment failure in the field are unclear (*53*). Molecular markers are widely used for monitoring changes in drug resistance mutations in malaria parasites (*54–56*) and for evaluating benzimidazole resistance in nematode parasites of veterinary importance (*57, 58*). The discovery of the genetic basis of resistance to another schistosome drug (oxamniquine) (*59*) now makes genetic surveys possible to evaluate oxamniquine resistance in schistosome populations (*52, 60*). Identification of *Sm.TRPM_PZQ_* as a critical determinant of PZQ response, and determination of key residues that can underlie PZQ-R, now makes molecular surveillance possible for *S. mansoni*. We examined variation in *Sm.TRPM_PZQ_* in 259 parasites collected from locations from across the geographical range of this parasite. We were unable to confirm mutations in any of the key residues that block PZQ binding identified in the mutagenesis studies by Park *et al.* (*21*). However, we identified a stop codon in a single parasite isolated from a rodent from Oman (*61*) indicating a low frequency of PZQ-R resistance alleles (1/502, frequency = 0.002). This stop codon was in heterozygous state so is unlikely to result in PZQ-R.

These results are extremely encouraging for control programs, but should be viewed with considerable caution for three reasons. First, the regulatory regions determining gene or isoform expression of *Sm.TRPM_PZQ_* are currently not known, so we were unable to examine regulatory variants of *Sm.TRPM_PZQ_* in this screen. Such variants could reduce expression of *Sm.TRPM_PZQ_* resulting in PZQ resistance. We note that coding variants underlying PZQ-R phenotype were not found in our laboratory SmLE-PZQ-ER parasites, suggesting that regulatory changes may underlie this trait. Second, *Sm.TRPM_PZQ_* is a large gene (120 kb and 41 exons) that is poorly captured by genome sequencing of field samples. We were able to successfully sequence 36/41 exons, including those that directly interact with PZQ (*21*), using exome capture methods (*51, 52*). However, improved sequence coverage will be needed for full length sequencing of this gene. Third, the parasite samples we examined did not come from hotspot regions where regular mass drug administration of PZQ has failed to reduce *S. mansoni* burdens (*35, 62*). Targeted sequencing of miracidia from these populations will be extremely valuable to determine if there are local elevations in *Sm.TRPM_PZQ_* variants, or if particular variants are enriched in parasites surviving PZQ treatment. Ideally, such sequence surveys should be partnered with functional validation studies in which variant *Sm.TRPM_PZQ_* are expressed in HEK293 cells to determine their response to PZQ exposure (*21*). As the homologous genes in *S. haematobium* and *S. japonicum* are likely to be activated by PZQ, this molecular approach to screening for PZQ resistance should be equally applicable to all major schistosome species infecting humans.

## MATERIALS AND METHODS

### Study design

This study was designed to determine the genetic basis of PZQ-R, and was stimulated by the initial observation that a laboratory *S. mansoni* population generated through selection with PZQ contained both PZQ-S and PZQ-R individuals. The project had 6 stages: (i) <u>QTL location</u>. We conducted a genome-wide association study (GWAS). This involved measuring the PZQ-response of individual worms, pooling those showing high levels of resistance and low levels of resistance, sequencing the pools to high read depth, and then identifying the genome regions showing significant differences in allele frequencies between high and low resistance parasites. (ii) <u>Fine mapping of candidate genes</u>. We identified potential candidate genes in these QTL regions, through examination of gene annotations, and exclusion of genes that are not expressed in adults. We also determined the mode of inheritance of the PZQ-R locus. (iii) <u>Marker assisted purification of PZQ-S and PZQ-R parasites</u>. To separate PZQ-R and PZQ-S parasites into “pure” populations, we genotyped larval parasites for genetic markers at the PZQ-R locus and infected rodents with genotypes associated with PZQ-R or PZQ-S. We then measured the IC_50_ for each of the purified populations to confirm their PZQ response. (iv) <u>Characterization of purified SmLE-PZQ-ER and SmLE-PZQ-ES populations</u>. Separation of SmLE-PZQ-ES and SmLE-PZQ-ER parasite populations allowed us to characterize these in more detail. Specifically, we measured expression in juvenile and adult worms of both sexes in both populations. We also quantified parasite fitness traits. (v) <u>Functional analysis</u>. We used RNAi and chemical manipulation approaches to modulate activity of candidate genes and determine the impact of PZQ-resistance. We also used transient expression of candidate genes in cultured mammalian cells, to determine the impact of particular SNPs on response to PZQ. (vi) <u>Survey of PZQ-resistance variants in field collected parasites</u>. Having determined the gene underlying PZQ-R in laboratory parasites, we examined sequence variation in this gene in a field collection of *S. mansoni* parasites to examine the frequency of sequence variants predicted to result in PZQ-resistance. Methods are described in detail (File S1) and in brief below.

### Ethics statement

This study was performed following the Guide for the Care and Use of Laboratory Animals of the National Institutes of Health. The protocol was approved by the IACUC of Texas Biomed (permit number: 1419-MA and 1420-MU). Ethical permission for collection of samples from humans are described in Chevalier *et al.* (*52*).

### Biomphalaria glabrata s*nails and* Schistosoma mansoni *parasites*

We used inbred albino *Biomphalaria glabrata* snails (line Bg121 (*63*)). The SmLE parasite population was originally obtained from an infected patient in Brazil (*64*). The SmLE-PZQ-R schistosome population was generated by applying a single round of PZQ selection pressure on SmLE parasites at both snail and rodent stages (*26*) and has been maintained in our laboratory since 2014.

### Drug resistance tests

#### Dose-response curves to PZQ in SmLE and SmLE-PZQ-R populations

We measured PZQ sensitivity by examining worm motility (*65*) in SmLE and SmLE-PZQ-R parasite populations. Ten adult males from SmLE or SmLE-PZQ-R populations were placed into each well of a 24-well microplate containing DMEM complete media (*59*). We performed control and experimental groups in triplicate (*N*=240 worms/population). We exposed adult worms to PZQ (0, 0.1 0.3, 0.9, 2.7, 8.1, 24.3 and 72.9 µg/mL) for 24h. Worms were washed (3x) in drug-free medium and incubated (37°C, 5% CO_2_) for 2 days. The parasites were observed daily for 5 days and the number of dead worms scored (i.e., opaque worms without movement).

#### Metabolic assessment of worm viability using L-lactate assay

We adapted a method for metabolic assessment of worm viability using an L-lactate assay (*44*). SmLE-PZQ-R adult males were individually placed in 96-well plates with filter insert (Millipore) in DMEM complete media. We added PZQ (24.3 µg/mL in DMEM complete media) while controls were treated with the drug diluent DMSO. At 48h post-treatment, the supernatant was collected from each well and immediately stored at −80 °C until processing. We measured L-lactate levels in the supernatants with a colorimetric L-lactate assay kit (Sigma).

### Genome wide association analysis and QTL mapping

#### Schistosome infections

We collected eggs from livers of hamsters infected with SmLE-PZQ-R (*66*) and exposed 1,000 Bg121 snails to 5 miracidia/snail. After 30 days, snails were individually exposed to light to shed cercariae. We exposed 8 hamsters to 840 cercariae (4 cercariae from each snail). We collected adult worms by hamster perfusion 45 days later.

#### Phenotypic selection

We plated SmLE-PZQ-R adult males individually in 96-well plates in DMEM complete media and treated with a 24.3 µg/mL PZQ. A control group was treated with the drug diluent DMSO. This experiment was done twice independently. A total of 590 and 691 adult males were collected, cultured *in vitro* and exposed to PZQ.

We collected media supernatants in 96-well PCR plates after 24h in culture (pre-treatment) and 48h post-treatment for quantifying L-lactate levels. We took the 20% of the treated worms releasing the highest amount of L-lactate (average L-lactate production ± SD: Experiment 1 = 61.44 nmol/h ± 13.16 / Experiment 2 = 56.38 nmol/h ± 10.82) and the 20% of the treated worms releasing the lowest amount of L-lactate (average L-lactate production ± SD: Experiment 1 = 28.61 nmol/h ± 5.32 / Experiment 2 = 23.4 nmol/h ± 4.14), 48h post PZQ treatment.

#### DNA extraction and library preparation

We sequenced whole genomes of the two pools of recovered (Experiment 1: 116 worms / Experiment 2: 137 worms) and susceptible worms (Experiment 1: 116 worms / Experiment 2: 137 worms) and measured allele frequencies in each pool to identify genome regions showing high differentiation. We extracted gDNA from pools (Blood and Tissue kit, Qiagen) and prepared whole genome libraries in triplicate (KAPA HyperPlus kit, KAPA Biosystems). Raw sequence data are available at the NCBI Sequence Read Archive (PRJNA699326).

#### Bioinformatic analysis

We used Jupyter notebook and scripts for processing the sequencing data and identifying the QTL (DOI: <u>10.5281/zenodo.5297220</u>).

##### a. Sequence analysis and variant calling

We aligned the sequencing data against the *S. mansoni* reference genome using BWA and SAMtools, used GATK to mark PCR duplicates and recalibrate base scores, the HaplotypeCaller module of GATK to call variants (SNP/indel) and GenotypeGVCFs module to perform joint genotyping. We merged VCF files using the MergeVcfs module. All steps were automated using Snakemake.

##### b. QTL identification

We examined the difference in allele frequencies between low and high L-lactate parasites across the genome by calculating a *Z*-score at each bi-allelic site, weighed *Z*-scores by including the number of worms in each treatment and the difference in the total read depth across the triplicated libraries of each treatment at the given variant, and combined *Z*-scores generated from each biological replicate. Bonferroni correction was calculated with α = 0.05.

### Relationship between worm genotype at chr. 2 and 3 and PZQ-R phenotype

We phenotyped and genotyped individual worms to validate the impact of worm genotypes on PZQ resistance phenotype and to determine the mode of inheritance of PZQ-R.

#### Measuring PZQ-R in individual worms

We plated 120 SmLE-PZQ-R adult males individually in 96-well plates, treated them with PZQ (24.3 µg/mL) and collected media supernatants pre- (24 h) and post- (48 h) treatment, and used L-lactate assays to determine PZQ-R status. gDNA was extracted from each worm individually using Chelex.

#### PCR-RFLP for chr.2 and chr.3 loci

We designed PCR-RFLP to genotype single worms at loci marking QTL peaks on chr. 2 (C>A, chr SM_V7_2: 1072148) and chr. 3 (T>C, chr SM_V7_3: 741987) (Table S4). To achieve this, we digested PCR amplicons for chr. 2 with BslI and chr. 3 with Mse1, which were visualized by 2% agarose gel electrophoresis.

#### Quantitative PCR of copy number variation (CNV) in single worms

We genotyped individual worms for a deletion on chr. 3 (position 1220683-1220861 bp) using a qPCR assay. Methods and primers are described in Table S4.

### Marker assisted selection of resistant and susceptible parasite populations

#### Selection of SmLE-PZQ-ER and SmLE-PZQ-ES populations

We separated the polymorphic SmLE-PZQ-R population based on chr. 3 QTL genotype using PCR-RFLP. We exposed 960 Bg121 snails to single SmLE-PZQ-R miracidia (*66*). We identified infected snails (*N*=272) 4 weeks later, collected cercariae from individual snails, extracted cercarial DNA (*66*), and genotyped each parasite at the chr. 3 locus (Homozygous R/R: n=89; Homozygous S/S: n=39; Heterozygous R/S: n=117), and determined their gender by PCR (*67*). We selected 32 R/R parasites and 32 S/S genotypes. For both R/R and S/S we used 13 males and 19 females. We exposed 5 hamsters to 800 cercariae of 32 R/R genotypes parasites and 5 hamsters to 800 cercariae 32 S/S genotyped parasites. This single generation marker assisted selection generated two subpopulations: SmLE-PZQ-ER enriched in parasites with R/R genotype, and SmLE-PZQ-ES enriched with S/S genotypes.

#### PZQ IC_50_ with SmLE-PZQ-ER and SmLE-PZQ-ES

Forty-five days after infection, we recovered adult worms of SmLE-PZQ-ER and SmLE-PZQ-ES subpopulations from hamsters. We placed adult males in 96-well plates in DMEM complete media. We determined PZQ dose-response using the same doses as for SmLE-PZQ-R.

#### gDNA extraction and library preparation and bioinformatics

We recovered the F1 SmLE-PZQ-ER and SmLE-PZQ-ES worms and extracted gDNA from pools of adult males or females separately and prepared whole genome libraries from these pools as described above. Sequence data are available at the NCBI Sequence Read Archive (accession numbers PRJNA701978). The analysis was identical to the GWAS and QTL mapping analysis.

### Transcriptomic analysis of resistant and susceptible schistosome worms at juvenile and adult stages

#### Sample collection

We recovered SmLE-PZQ-ER and SmLE-PZQ-ES worms from hamsters at 28 days (juveniles) or 45 days (adults) post-infection. For each subpopulation, we collected 3 biological replicates of 30 male or 30 female juvenile worms, and 3 biological replicates of 30 males or 60 female adult worms. Different numbers of males and females were used to provide sufficient amount of RNA material for library preparation.

#### RNA extraction and RNA-seq library preparation

##### a. RNA extraction

We extracted total RNA from juvenile and adult worm pools (RNeasy Mini kit, Qiagen), quantified the RNA recovered (Qubit RNA Assay Kit, Invitrogen) and assessed the RNA integrity by TapeStation (Agilent - RNA integrity numbers: ∼8.5–10 for all samples).

##### b. RNA-seq library preparation

We generated RNA-seq libraries using the KAPA Stranded mRNA-seq kit (KAPA Biosystems) from 500 ng RNA for each library and did 150 bp pair-end sequencing. Raw sequence data are available at the NCBI Sequence Read Archive (accession numbers PRJNA704646).

##### c. Bioinformatic analysis

To identify differentially expressed genes, we aligned the sequencing data against the *S. mansoni* reference genome using STAR. We quantified gene and isoform abundances by computing transcripts per million values using RSEM and compared abundances between groups (ES/ER, males/females, juveniles/adults) using the R package EBSeq. Jupyter notebooks and associated scripts are available on Zenodo (DOI: <u>10.5281/zenodo.5297218</u>)

### Manipulation of Sm.TRPM_PZQ_ channel expression or function

#### RNA interference

We attempted RNA interference to functionally validate the role of *Sm.TRPM_PZQ_* in PZQ resistance. This approach was unsuccessful. Detailed methodology is available in File S1 and Table S2.

#### Specific Sm.TRPM_PZQ_ chemical inhibitor and activator

We used specific chemical inhibitor and activator (Chulkov *et al.*, in prep) to manipulate the function of Sm.TRPM_PZQ_ to examine the impact on PZQ response. We placed individual SmLE-PZQ-ER and SmLE-PZQ-ES adult males in 96-well plates containing DMEM complete media. After 24h, worms from each population were treated either with a cocktail combining PZQ (1 µg/mL) and the Sm.TRPM_PZQ_ blocker (ANT1 at 50 µM) or the Sm.TRPM_PZQ_ activator (AG1 at 10, 25 or 50 µM) or the drug diluent (DMSO). We also set up control plates to evaluate the impact of Sm.TRPM_PZQ_ blocker or activator alone. Worms were exposed to these drug cocktails for 24h, washed 3 times with drug-free medium, and incubated (37°C, 5% CO_2_) for 2 days. We collected media supernatants before treatment and 48h post-treatment and quantified L-lactate.

### *In vivo* parasite survival after PZQ treatment

We exposed two groups of 12 female Balb/C mice to either SmLE-PZQ-ER or SmLE-PZQ-ES cercariae. Each mouse was infected by tail immersion using 80 cercariae from 40 infected snails. Forty days post-infection, we treated mice by oral gavage with either 120 mg/kg of PZQ (1% DMSO + vegetable oil in a total volume of 150 µL) or the same volume of drug diluent only (control group). We recovered worms 50 days post-infection.

### *Sm.TRPM_PZQ_* variants in *S. mansoni* field samples

#### Variants identification in exome sequence data from natural S. mansoni parasites

We utilized *S. mansoni* exome sequence data from Africa, South America and the middle East to investigate variation in *Sm.TRPM_PZQ_*. African miracidia were from the Schistosomiasis collection at the Natural History Museum (SCAN) (*68*), while Brazilian miracidia and Omani cercariae and adult worms were collected by laboratories at Texas Biomed or University of Perpignan, respectively (*52*). We have previously described exome sequencing methods for *S. mansoni* (*51, 52*). Data were analyzed as described in Chevalier et al. (*52*). The code is available in Jupyter notebook and associated scripts (DOI: <u>10.5281/zenodo.5297222</u>).

#### Sanger re-sequencing to confirm the presence of the Sm.TRPM_PZQ_ field variants

To confirm the presence of variants in *Sm.TRPM_PZQ_* when read depth was <10 reads, we performed Sanger re-sequencing of exons of interest (*21*). Primers and conditions are listed in Table S4. We scored variants using PolyPhred software (v6.18). Sanger traces are available on Zenodo (DOI: <u>10.5281/zenodo.5204523</u>). These analyses are available in the Jupyter notebook and associated scripts (DOI: <u>10.5281/zenodo.5297222</u>).

#### Functional analysis

We determined whether proteins produced by variant *Sm.TRPM_PZQ_* alleles were activated by PZQ following Park et al. (*21*).

### Statistical analysis

Statistical analyzes and graphs were performed using R software (v3.5.1). We used the drc package from R to analyze dose-response datasets and Readqpcr and Normqpcr packages to analyze RT-qPCR datasets. For non-normal data (Shapiro test, *p* < 0.05), we used Chi-square test or Kruskal-Wallis test followed by pairwise Wilcoxon-Mann-Whitney post-hoc test or a Wilcoxon-Mann-Whitney test. For normal data, we used one–way ANOVA or a pairwise comparison Welsh *t*-test. The confidence intervals were set to 95% and *p*-values < 0.05 were considered significant.

## Supporting information

Supplementary table 1 - Genes in QTL regions on chr 2 and 3.

Supplementary table 3 -Mutations present in Sm.TRPMPZQ in natural schistosome populations

## Acknowledgments

We thank the Vivarium of the Southwest National Primate Research Center (SNPRC) for providing rodent care. This research was supported by a Cowles fellowship (WLC) from Texas Biomedical Research Institute (13-1328.021), and NIH R01AI133749 (TJCA) and was conducted in facilities constructed with support from Research Facilities Improvement Program grant C06 RR013556 from the National Center for Research Resources. SNPRC research at Texas Biomedical Research Institute is supported by grant P51 OD011133 from the 570 Office of Research Infrastructure Programs, NIH. Miracidia samples for sequence analysis were provided by Schistosomiasis Collection at the Natural History Museum (SCAN) funded by the Wellcome Trust (grant 104958/Z/14/Z) and curated by Muriel Rabone. We thank Dr Oumar T. Diaw and Moumoudane M. Seye (Institut Sénégalais de Recherches Agricoles, Senegal) for the collections in Senegal (EU-CONTRAST project); Dr Anouk Gouvras (Global Schistosomiasis Alliance (GSA)), Mariama Lamine and the Réseau International Schistosomiases Environnemental Aménagement et Lutte (RISEAL) team (Niger) for the collections in Niger; Dr Anouk Gouvras (GSA) and the National Institute for Medical Research (NIMR, Mwanza Tanzania) team Dr Teckla Angelo, Honest Nagai, Boniface Emmanuel, John Igogote, Sarah Buddenborg and Reuben Jonathan for the collections in Tanzania as part of the Schistosomiasis Consortium for Operational Research and Evaluation (SCORE) project (https://score.uga.edu).

## Funding

National Institutes of Health grant <u>1R01AI123434</u> (TJCA, PTL)

National Institutes of Health grant 1R01AI133749 (TJCA)

National Institutes of Health grant NIH R01AI145871 (JSM)

Ministry of Health in Oman, the Sultan Qaboos University (grant no. IG/ MED/MICR/00/01)

French Ministry of Foreign Affairs (French Embassy in Oman) (grants nos. 402419B, 402415K and 339660F)

CNRS-Sciences de la Vie (grants no. 01N92/0745/1 and 02N60/1340)

CNRS-Direction des Relations internationales (grants no. 01N92/0745 and 02N60/1340)

PICS-CNRS no. 06249: FRANC-INCENSE

## Author contributions

Conceptualization: WLC, PTL, TJCA

Methodology: WLC, FDC, PTL, JM

Investigation: WLC, FDC, ACM, AS, RD, JM

Visualization: WLC, FDC

Field samples: HM, GM, MI, KAM, SAY

Funding acquisition: TJCA, PTL, JM, HM, GM

Project administration: TJCA, PTL

Supervision: TJCA, WLC, FDC, PTL, JM

Writing – original draft: TJCA, WLC, FDC

Writing – review & editing: TJCA, WLC, FDC, PTL, JM, HM, GM

## Data and materials availability

Phenotype, genotype and fitness data are available from: Zenodo – DOI: 10.5281/zenodo.5209300

Sequence data is available from: PRJNA699326, PRJNA701978, PRJNA704646

Sequence data from field collected *S. mansoni* is available from: PRJNA439266, PRJNA560070, PRJNA560069, PRJNA743359

Code is available on Zenodo (DOIs: <u>10.5281/zenodo.5297220</u>, <u>10.5281/zenodo.5297218</u>, <u>10.5281/zenodo.5297222</u>)

## Supplementary Materials

### File S1. EXPANDED MATERIALS AND METHODS

#### Study design

This study was designed to determine the genetic basis of PZQ-R, and was stimulated by the initial observation that a laboratory *S. mansoni* population generated through selection with PZQ contained both PZQ-S and PZQ-R individuals. The project had 6 stages:(i) <u>QTL location</u>. We conducted a genome-wide association study (GWAS). This involved measuring the PZQ-response of individual worms, pooling those showing high levels of resistance and low levels of resistance, sequencing the pools to high read depth, and then identifying the genome regions showing significant differences in allele frequencies between high and low resistance parasites. (ii) <u>Fine mapping of candidate genes</u>. We identified potential candidate genes in these QTL regions, through examination of gene annotations, and exclusion of genes that are not expressed in adults. We also determined whether the loci determining PZQ-R are inherited in a recessive, dominant or co-dominant manner (iii) <u>Marker assisted purification of PZQ-S and PZQ-R parasites</u>. To separate PZQ-R and PZQ-S parasites into “pure” populations, we genotyped larval parasites for genetic markers in the QTL regions and infected rodents with genotypes associated with PZQ-R or PZQ-S. To verify that this approach worked, we then measured the IC_50_ for each of the purified populations. (iv) <u>Characterization of purified SmLE-PZQ-ER and SmLE-PZQ-ES populations</u>. Separation of SmLE-PZQ-ES and SmLE-PZQ-ER parasite populations allowed us to characterize these in more detail. Specifically, we measured expression in juvenile and adult worms of both sexes in both SmLE-PZQ-ES and SmLE-PZQ-ER parasites. We also quantified parasite fitness traits. (v) <u>Functional analysis</u>. We used RNAi and chemical manipulation approaches to modulate activity of candidate genes and determine the impact of PZQ-resistance. We also used transient expression of candidate genes in cultured mammalian cells, to determine the impact of particular SNPs on response to PZQ-exposure. (vi) <u>Survey of PZQ-resistance variants in field collected parasites</u>. Having determined the gene underlying PZQ-R in laboratory parasites, we examined sequence variation in this gene in a field collection of *S. mansoni* parasites collected to examine the frequency of sequence variants predicted to result in PZQ-resistance.

#### Ethics statement

This study was performed in accordance with the Guide for the Care and Use of Laboratory Animals of the National Institutes of Health. The protocol was approved by the Institutional Animal Care and Use Committee of Texas Biomedical Research Institute (permit number: 1419-MA and 1420-MU). Details of ethical permission for collection of samples from humans are described in Chevalier et al. (*52*).

#### Biomphalaria glabrata s*nails and* Schistosoma mansoni *parasites*

Uninfected inbred albino *Biomphalaria glabrata* snails (line Bg121 derived from 13-16-R1 line (*63*)) were reared in 10-gallon aquaria containing aerated freshwater at 26-28 °C on a 12L-12D photocycle and fed *ad libitum* on green leaf lettuce. All snails used in this study had a shell diameter between 8 and 10 mm. We used inbred snails to minimize the impact of snail host genetic background on the parasite life history traits (*66*).

The SmLE schistosome (*Schistosoma mansoni*) population was originally obtained from an infected patient in Belo Horizonte (Minas Gerais, Brazil) in 1965 and has since been maintained in laboratory (*64*), using *B. glabrata* NMRI, inbred Bg36 and Bg121 population as intermediate snail host and Syrian golden hamsters (*Mesocricetus auratus*) as definitive hosts. The SmLE-PZQ-R schistosome population was generated in Brazil by applying a single round of PZQ selection pressure on SmLE parasites at both snail and rodent stages (*26*). The SmLE-PZQ-R population has been maintained in our laboratory using *B. glabrata* NMRI, Bg36 and Bg121 snail population and hamsters as the definitive host since 2014.

#### Drug resistance tests

##### Dose-response curves to PZQ in SmLE and SmLE-PZQ-R populations

Drug sensitivity to Praziquantel (PZQ) was initially measured using a modified protocol (*65*) in SmLE and SmLE-PZQ-R parasite populations. Ten adult males, recovered by perfusion from infected hamsters (t+ 45 days post-infection) (*69*) from SmLE or SmLE-PZQ-R population were placed into each well of a 24-well microplate containing 1 mL of High glucose DMEM supplemented with 15% heat-inactivated fetal bovine serum, 100 U/mL penicillin and 100μg/mL streptomycin (DMEM complete media). We performed control and experimental groups in triplicate (*N*=240 worms/parasite populations). We exposed adult worms to PZQ (0.1 µg/mL, 0.3 µg/mL, 0.9 µg/mL, 2.7 µg/mL, 8.1 µg/mL, 24.3 µg/mL and 72.9 µg/mL) for 24h. Worms were then washed three times in drug-free medium and incubated (37°C, 5% CO_2_) for 2 days. Control groups were exposed only to the drug diluent dimethyl sulfoxide (DMSO). The parasites were observed daily under a stereomicroscope for the 5 days of the experiment and the number of dead worms visually scored. Worms were defined as “dead” if they showed no movements and became opaque. We scored PZQ-resistance as a binary trait: parasites that recovered were classed as resistant, while parasites that failed to recover were classed as sensitive.

##### Metabolic assessment of worm viability using L-lactate assay

We adapted a method for metabolic assessment of worm viability using L-lactate assay (*44*). Briefly, adult male SmLE-PZQ-R worms recovered from infected hamsters were placed individually in 96-well plates containing 100 µm mesh filter insert (Millipore) in 250 µL DMEM complete media, and allowed to adapt for 24 h. We added PZQ (24.3 µg/mL in DMEM complete media) for the PZQ treated group, while controls were treated with the same volume of drug diluent DMSO. We also added a heat-killed worm control group: adult male worms were placed into a microfuge tube containing distilled water and heated in a dry bath (80°C, 15 min), and then plated in 96-well plate with 100 µm mesh insert. Drug resistance test was conducted as described above (see *Dose-response curves to PZQ in SmLE and SmLE-PZQ-R populations*). At 48h post-treatment, the supernatant (125 µL) was collected from each well and immediately stored at −80 °C until processing.

We measured L-lactate levels in the supernatants of *in vitro* treated adult male worms with a colorimetric L-lactate assay kit (Sigma) using 96-well, optical clear-bottom plates (Corning) following the manufacturer’s specifications, with minor modifications. Briefly, 5 µL of supernatant were diluted into 20 µL of ddH_2_O to fit within the linear range of the assay. We then combine 24 µL of the assay buffer to 1 µL of diluted supernatant (1/5 dilution) in each test well and added 25 µL of the lactate reaction mix (24 µL of the assay buffer, 0.5 µL of enzyme mix and 0.5 µL of lactate assay probe - V_total_= 50 µL/well). We also made a L-lactate standard curve to allow accurate L-lactate quantification in worm supernatants. After 45 min of incubation in the dark at room temperature, the plate was read by a spectrophotometer (Molecular Devices) at 570 nm. Lactate quantities in worm supernatant were assessed following the manufacturer’s instruction, taking in account our dilution factor. All measurement series included a DMEM complete media control to determine the background lactate level, which was then subtracted from the L-lactate quantity of the respective measurements.

#### Genome wide association analysis and QTL mapping

##### Schistosome infections

Eggs were collected from livers of hamster infected with SmLE-PZQ-R and hatched under light for 30 min in freshwater to obtain miracidia (*66*). We then exposed one thousand Bg121 snails to five miracidia/snail. After 30 days, snails were individually exposed to light in 24-well plates to shed cercariae. Eight hamsters were exposed to 840 cercariae (4 cercariae/snail) from a batch of 210 shedding snails. We euthanized hamsters after 45 days to collect adult worms.

##### Phenotypic selection

Adult SmLE-PZQ-R worms were collected, separated by sex and males were plated individually in 96-well plates (60 worms per plate) containing 100uM mesh filter insert (Millipore) in 250 µL of DMEM complete media and treated with a dose of 24.3 µg/mL PZQ as describe above (i.e. *Metabolic assessment of worms viability using L-lactate assay*). A group of 12 worms were treated with the same volume of drug diluent DMSO. This GWAS experiment was done twice independently. A total of 590 and 691 adult male worms were collected, cultured *in vitro* and exposed to PZQ for the two experiments respectively.

Worm media supernatants (125 µL) were collected in 96-well PCR plates after 24h in culture (to assess the viability of the worms before PZQ treatment – adult male worms should release ≥ 40 nmol/h of L-lactate in supernatant) and 48h post-treatment (to assess their viability after PZQ treatment). Plates containing supernatant were immediately stored at −80 °C until processing. Lactate levels in supernatants were quantified as described above (see *Metabolic assessment of worm viability using L-lactate assay*). We phenotype the worms and categorize them into two groups: i) Recovered worms (i.e. releasing ≥ 40 nmol/h of L-lactate in supernatant) and ii) Susceptible worms (i.e. releasing less 40 nmol/h of L-lactate in supernatant). Among these two groups, we took the 20% of the treated worms releasing in their media the highest amount of L-lactate (average L-lactate production ± SD: Experiment 1 = 61.44 nmol/h ± 13.16 / Experiment 2 = 56.38 nmol/h ± 10.82) and the 20% of the treated worms releasing the lowest amount of L-lactate (average L-lactate production ± SD: Experiment 1 = 28.61 nmol/h ± 5.32 / Experiment 2 = 23.4 nmol/h ± 4.14), 48h post PZQ treatment respectively.

##### DNA extraction and library preparation

We sequenced whole genomes of the two pools of recovered (i.e. resistant to PZQ, Experiment 1: 116 worms / Experiment 2: 137 worms) and susceptible worms (Experiment 1: 116 worms / Experiment 2: 137 worms). We then estimated allele frequencies in each pool to identify genome regions showing high differentiation.

a. ***gDNA extraction:*** We extracted gDNA from pools of worms using the Blood and Tissue kit (Qiagen). We homogenized worms in DNA extraction kit lysis buffer using sterile micro pestles., incubated homogenates (56 °C, 2 hour) and recovered gDNA in 200 µL of elution buffer. We quantified the worm gDNA recovered using the Qubit dsDNA HS Assay Kit (Invitrogen).
b. ***Whole genome library preparation and sequencing:*** We prepared whole genome libraries from pools of worm gDNA in triplicate using the KAPA HyperPlus kit (KAPA Biosystems) according to the manufacturer’s protocol. For each library, we sheared 100 ng of gDNA by adaptive focused acoustics (Duty factor: 10%; Peak Incident Power: 175; Cycles per Burst: 200; Duration: 180 seconds) in AFA tubes (Covaris S220 with SonoLab software version 7 (Covaris, Inc., USA)) to recover fragmented DNA (150-200 bp). Library indexing was done using KAPA Dual Adapters at 15 µM for 1h. We used 6 PCR cycles for post-ligation library amplification. We performed size selection on the indexed-amplified libraries using KAPA Pure bead (0.7x first upper size cut; 0.9x second lower size cut). We quantified libraries by qPCR using KAPA library quantification kit (KAPA Biosystems) and their respective fragment size distribution was assessed by TapeStation (Agilent). We sequenced the libraries on a HiSeq X sequencer (Illumina) using 150 bp pair-end reads. Raw sequence data are available at the NCBI Sequence Read Archive (PRJNA699326).

##### Bioinformatic analysis

Jupyter notebook and scripts used for processing the sequencing data and identifying the QTL are available on Zenodo (DOI: <u>10.5281/zenodo.5297220</u>).

a. ***Sequence analysis and variant calling:*** We aligned the sequencing data against the *S. mansoni* reference genome (schistosoma_mansoni.PRJEA36577.WBPS14) using BWA (v0.7.17) and SAMtools (v1.10). We used GATK (v4.1.8.1) to mark PCR duplicates and recalibrate base scores. We used the HaplotypeCaller module of GATK to call variants (SNP/indel) and the GenotypeGVCFs module to perform a joint genotyping on each chromosome or unassembled scaffolds. We merged VCF files using the MergeVcfs module. All these steps were automatized using Snakemake (v5.14.0).
b. ***QTL identification:*** We expect the genome region underlying resistance to be enriched in variants from high L-lactate producing worms. To evaluate statistical evidence for such enrichment, we examined the difference in allele frequencies between low and high L-lactate parasites across the genome by calculating a *Z*-score at each bi-allelic site. To minimize bias, we weighed *Z*-scores by including the number of worms in each treatment and the difference in the total read depth across the triplicated libraries of each treatment at the given variant. We calculated *Z*-scores for each biological replicate as follows:

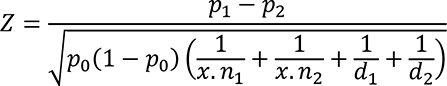

where *p*1 and *p*2 are the estimated allele frequencies in the low and high L-lactate parasites pools, respectively; *p*0 is the allele frequency under the null hypothesis H0: *p*1 = *p*2 estimated from the average of *p*1 and *p*2; *n*1 and *n*2 are the number of worms in the low and high L-lactate parasites pools, respectively, factor *x* for each *n* reflecting the ploidy state (*x*=2); and *d*1 and *d*2 are the sequencing depths for the low and high L-lactate parasite pools, respectively. We combined *Z*-scores generated from each biological replicate as follows:

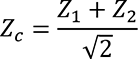

where *Z*_1_ and *Z*_2_ were *Z*-scores from replicate 1 and 2, respectively. The p-values were obtained by comparing the negative absolute value of *Z*-scores to the standard normal distribution. To determine the significant threshold, Bonferroni correction was calculated with α = 0.05. These analyses are available in the Jupyter notebook and associated scripts (DOI: <u>10.5281/zenodo.5297220</u>).

#### Relationship between worm genotype at chr. 2 and 3 and PZQ-R phenotype

To validate the impact of worm genotypes on its PZQ resistance phenotypes and determine whether PZQ-R shows recessive, dominant or codominant inheritance, we determined the PZQ-R phenotype of individual worms, which were then genotyped for markers at the peak of the QTLs located.

##### Measuring PZQ-R in individual worms

We collected 120 SmLE-PZQ-R adult male worms, plated them individually in 96-well plates containing a mesh filter insert, cultured them *in vitro*, treated them with PZQ (24.3 µg/mL) and collected media supernatants before (after 24h in culture) and 48h post-treatment, and used L-lactate assays to determine PZQ-R status (see *Phenotypic selection*). We extracted gDNA from each worm individually. Briefly, we transferred worms into 96-well PCR plates, added 100 µL of 6% Chelex® solution containing 1% Proteinase K (20 mg/mL), incubated for 2h at 56 °C and 8 min at 100 °C, and transferred the supernatant containing worm gDNA into fresh labeled 96-well plates.

##### PCR-RFLP conditions for chr.2 and chr.3 loci

We used PCR-RFLP to genotype single worms at loci marking QTL peaks on chr. 2 (C>A, chr SMV7_2: 1072148) and chr. 3 (T>C, chr SMV7_3: 741987). Primers were designed using PerlPrimer v1.21.1 (Table S4). We digested PCR amplicons for chr. 2 with BslI (NEB) and chr. 3 with Mse1 (NEB), and visualized digested PCR amplicons by 2% agarose gel electrophoresis.

##### Quantitative PCR validation of copy number variation (CNV) in single worms

We genotyped each individual worm for a deletion identified on chr. 3 at position 1220683-1220861 bp using a custom quantitative PCR assay. This was done to examine the association between deletion of this genomic region and PZQ resistant genotype. We quantified the copy number in this region relative to a single copy gene from *S. mansoni* (α-tubulin 2) (*66*). The CNV genotype for each parasite corresponds to the ratio of the CNV copy number and the α-tubulin 2 gene copy number: 0=complete deletion, 0.5=one copy, 1=two copies. Methods and primers are described in Table S4. We then compared individual worm phenotypes for each of the three CNV genotypes (0, 1 or 2 copies) to determine the association between CNV and PZQ response.

#### Marker assisted selection of resistant and susceptible parasite populations

##### Selection of SmLE-PZQ-ER and SmLE-PZQ-ES populations

We separated the polymorphic SmLE-PZQ-R schistosome population based on chr. 3 QTL genotype using the PCR-RFLP as described. We exposed 960 inbred *B. glabrata* Bg121 snails to one miracidium SmLE-PZQ-R (*66*). At four weeks post-exposure, we identified infected snail (*N*=272), and collected cercariae from individual snails. We extracted cercarial DNA using 6%Chelex (*66*)), and genotyped each parasite for chr. 3 locus using our PCR-RFLP (Homozygous R/R: n=89 – 36%; Homozygous S/S: n=39 – 16%; Heterozygous R/S: n=117 – 49%) and determine their gender by PCR (*67*). We selected 32 R/R parasites (homozygous for the *Sm.TRPM_PZQ_* resistant-associated allele) and 32 S/S genotypes (i.e. homozygous for the *Sm.TRPM_PZQ_* sensitive-associated allele). For both R/R and S/S we used 13 males and 19 females. We exposed 5 hamsters to 800 cercariae of 32 R/R genotypes parasites and 5 hamsters to 800 cercariae 32 S/S genotyped parasites. This single generation marker assisted selection procedure generates two subpopulations: SmLE-PZQ-ER is expected to be enriched in parasites with R/R genotype and to show strong PZQ-R, while SmLE-PZQ-ES is enriched for S/S genotypes and is expected to be highly sensitive to PZQ).

##### PZQ IC_50_ with SmLE-PZQ-ER and SmLE-PZQ-ES

Forty-five days after exposure to cercariae, we euthanized and perfused hamsters to recover adult schistosome worms from each subpopulation (SmLE-PZQ-ER and SmLE-PZQ-ES). We separated worms by sex and we set adult males in 96-well plates containing 100 µm mesh filter insert (Millipore) and cultured in 250 µL DMEM complete media as described.

We determined PZQ dose-response for both SmLE-PZQ-ER and SmLE-PZQ-ES population. We exposed individual worms (*N*=60/population/treatment) to PZQ (0.1 µg/mL, 0.3 µg/mL, 0.9 µg/mL, 2.7 µg/mL, 8.1 µg/mL, 24.3 µg/mL and 72.9 µg/mL) or drug diluent (DMSO control). Worm media supernatants (125 µL) were collected in 96-well PCR plates before treatment (after 24h in culture) and 48h post-treatment. We quantified L-lactate levels in supernatants described and we assess variation in L-lactate production for each individual worm.

##### gDNA extraction and library preparation

SmLE-PZQ-ER and SmLE-PZQ-ES parasite populations were maintained in our laboratory. We recovered the F1 worms from each populations and extract gDNA from pools of adult males (100 worms/population) and females (25 worms/population) separately as described above. Post extraction cleaning using Genomic DNA Clean & Concentrator (Zymo) have been performed on female qDNA pools to remove hemoglobin contaminants. We prepared whole genome libraries from these pools in triplicate using the KAPA HyperPlus kit (KAPA Biosystems) as described (see *Whole genome library preparation and sequencing*). We sequenced the libraries on a HiSeq X sequencer (Illumina) using 150 bp paired-end reads. Sequence data are available at the NCBI Sequence Read Archive (accession numbers PRJNA701978).

##### Bioinformatic analysis

The analysis was identical to the GWAS and QTL mapping analysis. This can be replicated with the Jupyter notebook and associated scripts (DOI: <u>10.5281/zenodo.5297220</u>).

#### Fitness of SmLE-PZQ-ER and SmLE-PZQ-ES parasite populations

The SmLE-PZQ-ER and SmLE-PZQ-ES populations of parasites has been maintained in our laboratory using B. glabrata Bg36 and Bg121 snail population and hamster definitive hosts for 12 generations. For each cycle and population, we exposed between 96 to 144 snails with 5-7 miracidia and two hamsters with 700 cercariae. Each generation, we measured snail survival (snails alive/the number of snails exposed) and snail infectivity (snail producing cercariae/number of snails tested for shedding) 4 weeks after miracidial exposure (*66*). We measured to hamsters for each population of parasites (worms recovered/number of cercariae used to infect each hamster).

#### Transcriptomic analysis of resistant and susceptible schistosome worms to PZQ at juvenile (28 days) and adult (45 days) stages

##### Sample collection

Juvenile and adult *S. mansoni* SmLE-PZQ-ER and SmLE-PZQ-ES worms were recovered by perfusion from hamsters at 28 days (juveniles) or 45 days (adults) post-infection. Worms from each population were placed in DMEM complete media, separated by sex, and aliquoted in sterile RNAse free microtubes which were immediately snap-frozen in liquid nitrogen and stored at −80 °C until RNA extractions. For each subpopulation, (SmLE-PZQ-ER or SmLE-PZQ-ES), we collected 3 biological replicates of 30 males and 3 replicates of 30 females for the 28d juvenile worms and 3 biological replicates of 30 males and 3 replicates of 60 females for the 45d adult worms.

##### RNA extraction and RNA-seq library preparation

###### a. RNA extraction

We extracted total RNA from all the *S. mansoni* adult and juvenile worms collected using the RNeasy Mini kit (Qiagen). Samples were randomized prior to RNA extraction to minimize batch effects. We homogenized worms in 600 µL RNA lysis buffer (RTL buffer, Qiagen) using sterile micro pestles, followed by passing the worm lysate 10 times through a sterile needle (23 gauge). We recovered total RNA in 25 µL elution buffer. We quantified the RNA recovered using the Qubit RNA Assay Kit (Invitrogen) and assessed the RNA integrity by TapeStation (Agilent - RNA integrity numbers of ∼8.5–10 for all the samples).

###### b. RNA-seq library preparation

We prepared RNA-seq libraries using the KAPA Stranded mRNA-seq kit (KAPA Biosystems) using 500ng RNA diluted in 50uL Tris-HCl (pH 8.0) for each library. We fragmented mRNA (6 min 94°C), indexed libraries using 3’-dTMP adapters (7 µM, 1 hour at 20°C), and used 6 PCR cycles for post-ligation library amplification. We quantified indexed libraries by qPCR (KAPA library quantification kit (KAPA Biosystems)) and assessed their fragment size distribution by TapeStation (Agilent). We sequenced the libraries on a HiSeq 4000 sequencer (Illumina) using 150 bp pair-end reads, pooling 12 RNA-seq libraries/lane. Raw sequence data are available at the NCBI Sequence Read Archive under accession numbers PRJNA704646.

###### c. Bioinformatic analysis

To identify differentially expressed genes between the different groups, we aligned the sequencing data against the *S. mansoni* reference genome (schistosoma_mansoni.PRJEA36577.WBPS14) using STAR (v2.7.3a). We quantified gene and isoform abundances by computing transcripts per million values using RSEM (v1.3.3). We compared these abundances between groups (ES/ER, males/females, juveniles/adults) using the R package EBSeq (v1.24.0). Jupyter notebooks and associated scripts are available on Zenodo (DOI: <u>10.5281/zenodo.5297218</u>).

#### Manipulation of Sm.TRPM_PZQ_ channel expression or function

##### RNA interference

We used RNA interference to knock down the expression of Smp_246790 gene in order to functionally validate the implication of *Sm.TRPM_PZQ_* on schistosome PZQ resistance. SmLE-PZQ-R adult male worms were freshly recovered from infected hamsters and placed in 24-well plates for in vitro culture (10 adult male worms/well).

###### a. siRNA treatment on S. mansoni adult male worms

Small inhibitory RNAs (siRNAs) targeting specific schistosome genes were designed using the on-line IDT RNAi Design Tool (https://www.idtdna.com/Scitools/Applications/RNAi/RNAi.aspx) (Table S2) and synthesized commercially by Integrated DNA Technologies (IDT, Coralville, IA). To deliver the siRNAs, we electroporated schistosome parasites (10 adults/group – each group done in triplicate) in 100 μL RPMI medium containing 2.5 μg siRNA or the equivalent volume of ddH_2_O (no siRNA control), in a 4 mm cuvette by applying a square wave with a single 20 ms impulse, at 125 V and at room temperature (Gene Pulser Xcell Total System (BioRad)) (*70*). Parasites were then transferred to 1 mL of complete DMEM media in 24-well plates. After overnight culture, medium was replaced with fresh DMEM complete media. We measured gene expression by quantitative real-time PCR (RT-qPCR) 2 days after siRNA treatment.

###### b. dsRNA treatment on S. mansoni adult male worms

We synthetized double-stranded RNA according to Wang et al. (*71*) (Table S2). For dsRNA treatment, 10 adult male worms/group (each group done in triplicate) were cultured in 1 mL DMEM complete media and treated with 90 ug dsRNA at day 0, day 1 and day 2. Media was changed every 24h and fresh dsRNA was added. On day 3, we harvested worms and measured gene expression by quantitative real-time PCR (RT-qPCR).

###### c. RNA extraction and gene expression analysis by RT-qPCR

We extracted total RNA from parasites (*N*=10 worms/sample) using the RNeasy Mini kit (Qiagen) (see *RNA extraction*). Complementary DNA (cDNA) was generated from extracted RNA (500 ng - 1 µg) using SuperScript-III and Oligo-dT primers (ThermoFisher). Relative quantification of genes of interest was performed in duplicate by qPCR analysis using QuantStudio 5 System (Applied Biosystems) and SYBR Green master mix (ThermoFisher), compared with a serially diluted standard of PCR products (generated from cDNA) for each of the gene tested (*66*). Standard curves allow evaluating the efficiency of each pairs of primers, for both housekeeping and target genes using QuantStudio Design and Analysis Software. Expression was normalized to SmGAPDH housekeeping gene (Table S2) using the efficiency E^ΔΔCt^ method (*72*).

##### Specific Sm.TRPM_PZQ_ chemical inhibitor and activator

We used specific chemical inhibitor and activators (Chulkov *et al.*, in prep) to manipulate the function of Sm.TRPM_PZQ_ to examine the impact on PZQ-response. We placed individual SmLE-PZQ-ER and SmLE-PZQ-ES adult male worms in 96-well plates with 100 µm mesh filter insert containing DMEM complete media and cultured described above (see *Metabolic assessment of worm viability using L-lactate assay*). After 24h in culture, 20 worms from each population were treated either with a cocktail combining PZQ (1 µg/mL) and i) 50 µM of Sm.TRPM_PZQ_ blocker (ANT1 MB2) or ii) 10 µM, 25 µM or 50 µM of Sm.TRPM_PZQ_ activator (AG1 MV1) respectively or iii) drug diluent (DMSO). We also set up control plates to evaluate the impact of Sm.TRPM_PZQ_ blocker or activator alone. In that case, 20 worms from each population were treated with a cocktail combining drug diluent DMSO and Sm.TRPM_PZQ_ blocker (MB2) or activator (MV1) at the same concentrations mentioned above. Worms were exposed to these drug cocktails for 24h, washed 3 times with drug-free medium, and incubated (37°C, 5% CO_2_) for 2 days.

We collected Worm media supernatants (125 µL) in 96-well PCR plates before treatment (after 24h in culture) and 48h post-treatment and L-lactate levels in supernatants were quantified as described above (see *Metabolic assessment of worm viability using L-lactate assay*). We used these results to determine the impact of blockers or activators on variation in L-lactate production.

#### *In vivo* parasite survival after PZQ treatment

We used 24 female Balb/C mice (purchased from Envigo at 6 weeks-old and housed in our facility for one week before use) split into two groups. Each group were exposed by tail immersion to to cercariae from SmLE-PZQ-ER (80 cercariae/mouse from 40 infected snails) or SmLE-PZQ-ES (80 cercariae/mouse from 40 infected snails) population. Each mouse was identified by a unique tattoo ID and an ear punch for assessing treatment status (PZQ or drug diluent control). Immediately after infection, we stained the content of each infection vial with 10 µL 0.4% Trypan blue and counted all the cercarial tails/heads or complete cercariae to determine the cercarial penetration rate for each mouse. We kept infected mice in 4 cages (2 cages/parasite populations and 6 animals per cage) at 21-22° C and 39%–50% humidity and monitored them daily.

Forty days post-infection, we weighed mice and treated them by oral gavage with either 120 mg/kg of PZQ (diluted in 1% DMSO + vegetable oil – Total volume given/mouse: 150 µL) or the same volume of drug diluent only (control group). To minimize batch effects, 3 mice were treated with PZQ and 3 with the drug diluent per cage for each parasite group (SmLE-PZQ-ER or SmLE-PZQ-ES). Mice were monitored daily until euthanasia and perfusion (*69*), at day 50 post-infection. We recorded the weight of each mouse before euthanasia. After euthanasia and perfusion, we also weighted the liver and spleen of each individual. We carefully recovered worms from the portal vein, liver and intestine mesenteric venules of each mouse. Worms were separated by sex and counted.

#### *Sm.TRPM_PZQ_* variants in *S. mansoni* field samples

##### Variants identification in exome-sequenced data from natural S. mansoni parasites

We utilized exome sequence data from *S. mansoni* from Africa, South America and the middle East to investigate variation in *Sm.TRPM_PZQ_*. African miracidia were from the Schistosomiasis collection at the Natural History Museum (SCAN) (*68*), Brazilian miracidia and Omani cercariae and adult worms were collected previously. We have previously described methods and generation of exome sequence from *S. mansoni* samples (*51, 52*). Data were analyzed the same way as described in Chevalier et al. (*52*). Code is available in Jupyter notebook and scripts (DOI: <u>10.5281/zenodo.5297222</u>).

##### Sanger re-sequencing to confirm the presence of the Sm.TRPM_PZQ_ field variants

To confirm the presence of the variants in *Sm.TRPM_PZQ_* gene from our exome-sequenced natural *S. mansoni* parasites (when read depth was <10 reads), we performed Sanger re-sequencing of eight *Sm.TRPM_PZQ_* exons (i.e. exon 3, 4, 23, 25, 27, 29 and 34) where either nonsense mutations (leading to truncated protein) or non-synonymous mutation located close to the PZQ binding site (*21*) were identified. Primers and conditions are listed in Table S4.

##### Sanger sequencing analysis

We scored variants using PolyPhred software (v6.18) (Nickerson et al., 1997) which relies on Phred (v0.020425.c), Phrap (v0.990319), and Consed (v29.0) software, analyzing each exon independently. We identified single nucleotide polymorphisms using a minimum phred quality score (-q) of 25, a minimum genotype score (-score) of 70, and a reference sequence of the *Sm.TRPM_PZQ_* gene from the chromosome 3 of *S. mansoni* reference genome (schistosoma_mansoni.PRJEA36577.WBPS14). Variant sites were labeled as non-reference alleles if they differed from the reference sequence. We identified insertion/deletion (indel) polymorphisms using a minimum phred quality score (-q) of 25, a minimum genotype score (-iscore) of 70. Polymorphisms were visually validated using Consed. Sanger traces are available on Zenodo (DOI: <u>10.5281/zenodo.5204523</u>). These analyses are available in the Jupyter notebook and associated scripts (DOI: <u>10.5281/zenodo.5297222</u>).

##### Functional analysis

We determined whether proteins produced by variant *Sm.TRPM_PZQ_* alleles were activated by PZQ following Park et al. (*21*) methods. In brief, we transiently expressed *Sm.TRPM_PZQ_* in HEK293 hamster cells, exposed transfected cells to PZQ, and measured Ca^2+^ influx to determine activation

#### Statistical analysis

All statistical analyzes and graphs were performed using R software (v3.5.1). We used the drc package from R to analyze all our dose-response datasets using a four-parameter log-logistic function to fit curves. We used the Readqpcr and Normqpcr packages to analyze all our RT-qPCR datasets, using the *efficiency*^ΔΔCt^ method. When data were not normally distributed (Shapiro test, *p* < 0.05), we compared results with non-parametric tests: Chi-square test, Kruskal-Wallis test followed by pairwise Wilcoxon-Mann-Whitney post-hoc test or a simple pairwise comparison Wilcoxon-Mann-Whitney test. When data followed a normal distribution, we used one–way ANOVA or a pairwise comparison Welsh *t*-test. The confidence interval of significance was set to 95% and *p*-values less than 0.05 were considered significant.

## SUPPLEMENTARY FIGURES

**Fig. S1.**
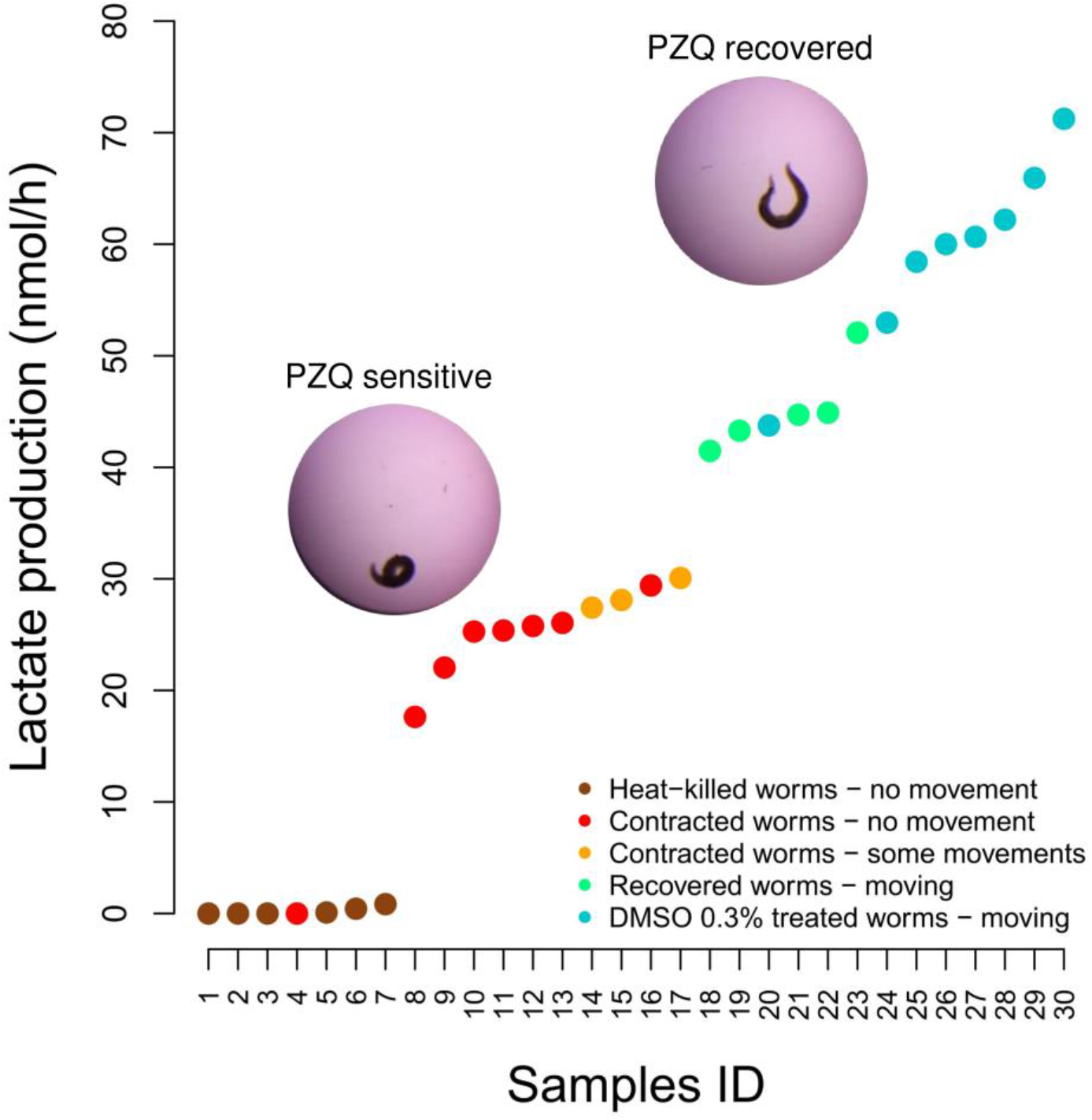
Development of a L-lactate assay for assaying worm recovery. Validation of the L-lactate assay for single male worms and correlation with their microscopic appearance and ability to regain movement after PZQ treatment (24.3 μg/mL). PZQ-S worms (contracted), that remain contracted after PZQ treatment produce significantly less amount of L-lactate released in the media compared to PZQ-R (recovered and motile) worms (Wilcoxon test, p = 0.0015; *N* = 30 worms).

**Fig. S2.**
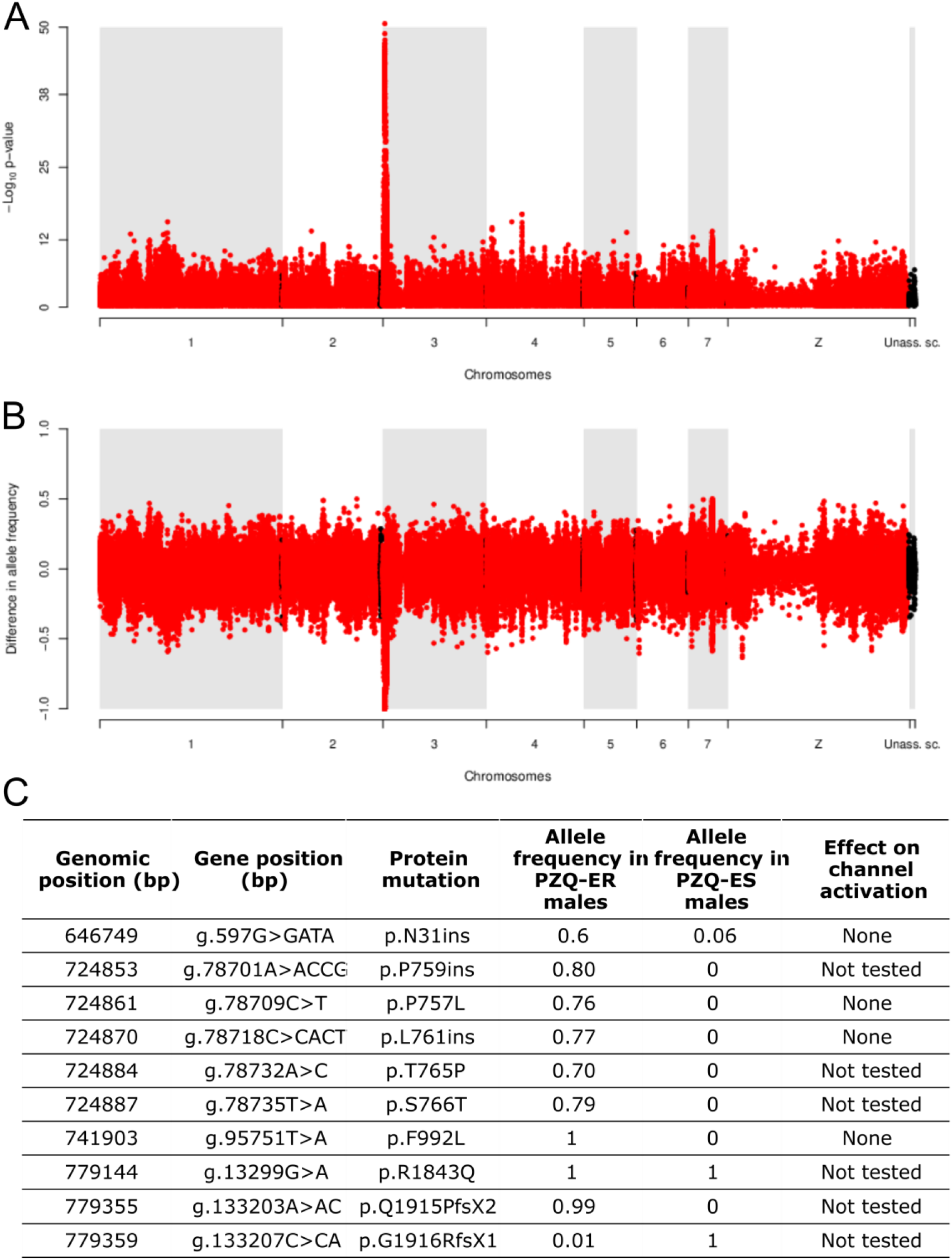
Validation of marker-assisted selection of SmLE-PZQ-ER and ES using Next Generation Sequencing (NGS). SmLE-PZQ-ER and ES differed only at the locus linked to PZQ resistance **(A)** Alternatively fixed allele were fixed for alternative alleles at the *Sm.TRPM_PZQ_*-741987C SNP genotyped (**B**; 1: fixed in ES, -1 fixed in ER), but showed similar allele frequencies across the rest of the genome. Segregating mutations between ER and ES are shown in table **(C)** We determined whether proteins expressing these variations were activated by PZQ following Park *et al.* methods (*21*).

**Fig. S3.**
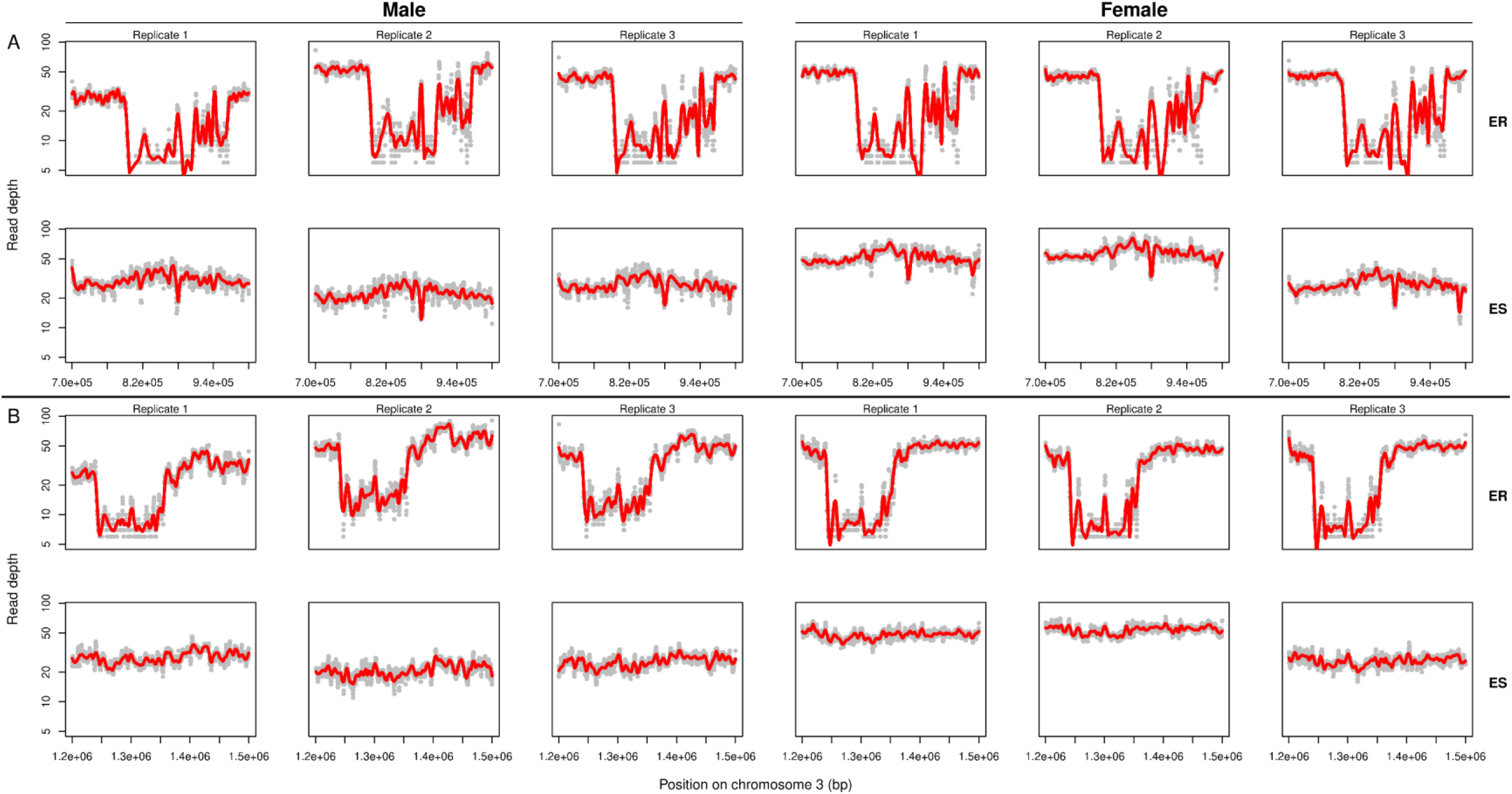
Large deletions adjacent to *Sm.TRPMPZQ* and SOX13 transcription factor. Sequencing of adult parasites recovered from SmLE-PZQ-ER and SmLE-PZQ-ES populations also revealed that the 100 kb **(A)** and the 150 kb **(B)** deletions were close to fixation in the SmLE-PZQ-ER population. The first row of each panel refers to the ER population, the second to the ES population. The first three columns refer to the female, the last three columns to the males.

**Fig. S4.**
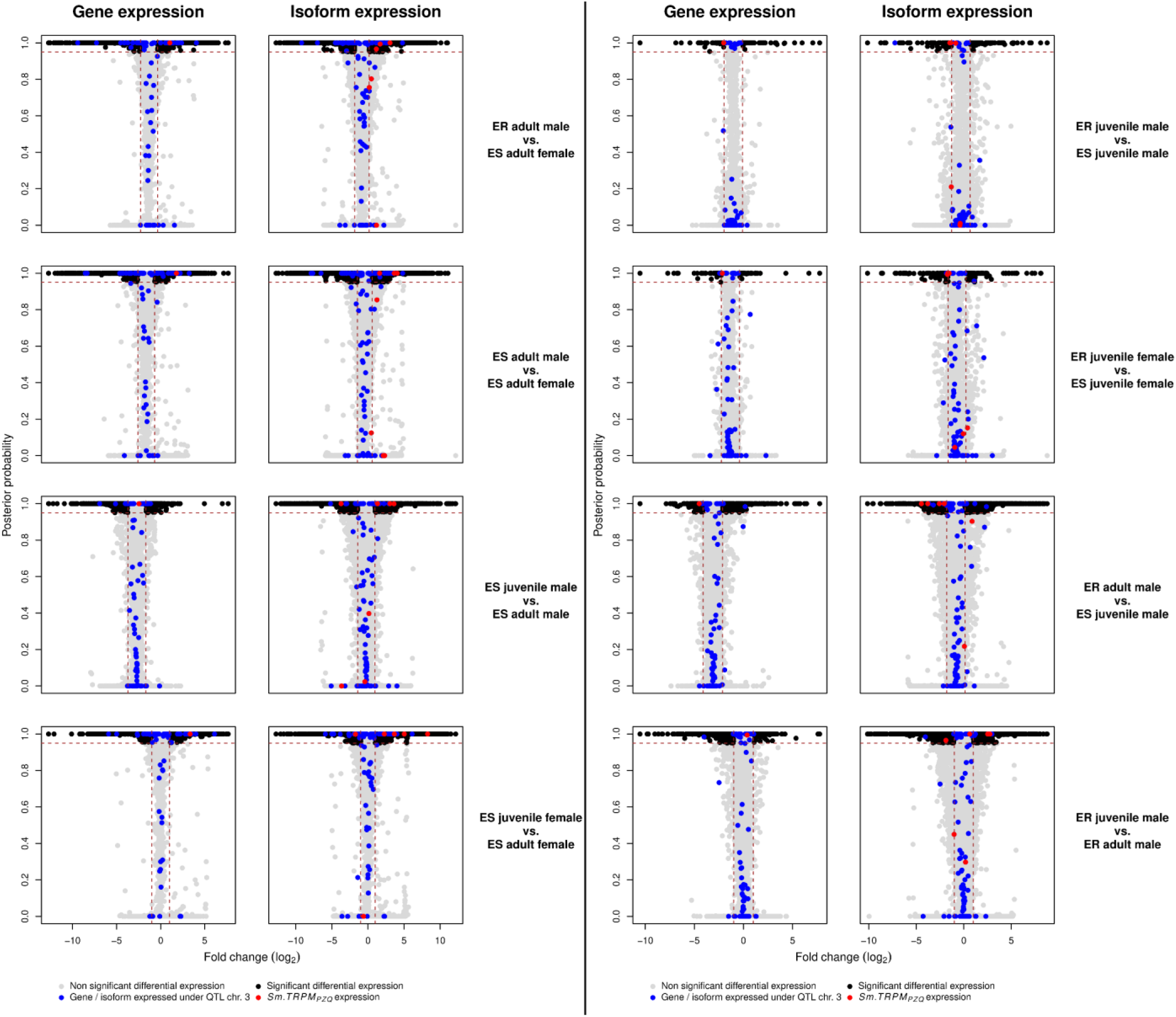
Detailed genes and isoforms expression in SmLE-PZQ-ER and SmLE-PZQ-ES parasites. Comparison of genes and isoforms expression between SmLE-PZQ-ER and ES parasites for each sex (i.e. male and female) and each parasite stage (i.e. adult worm and juvenile worm)

**Fig. S5.**
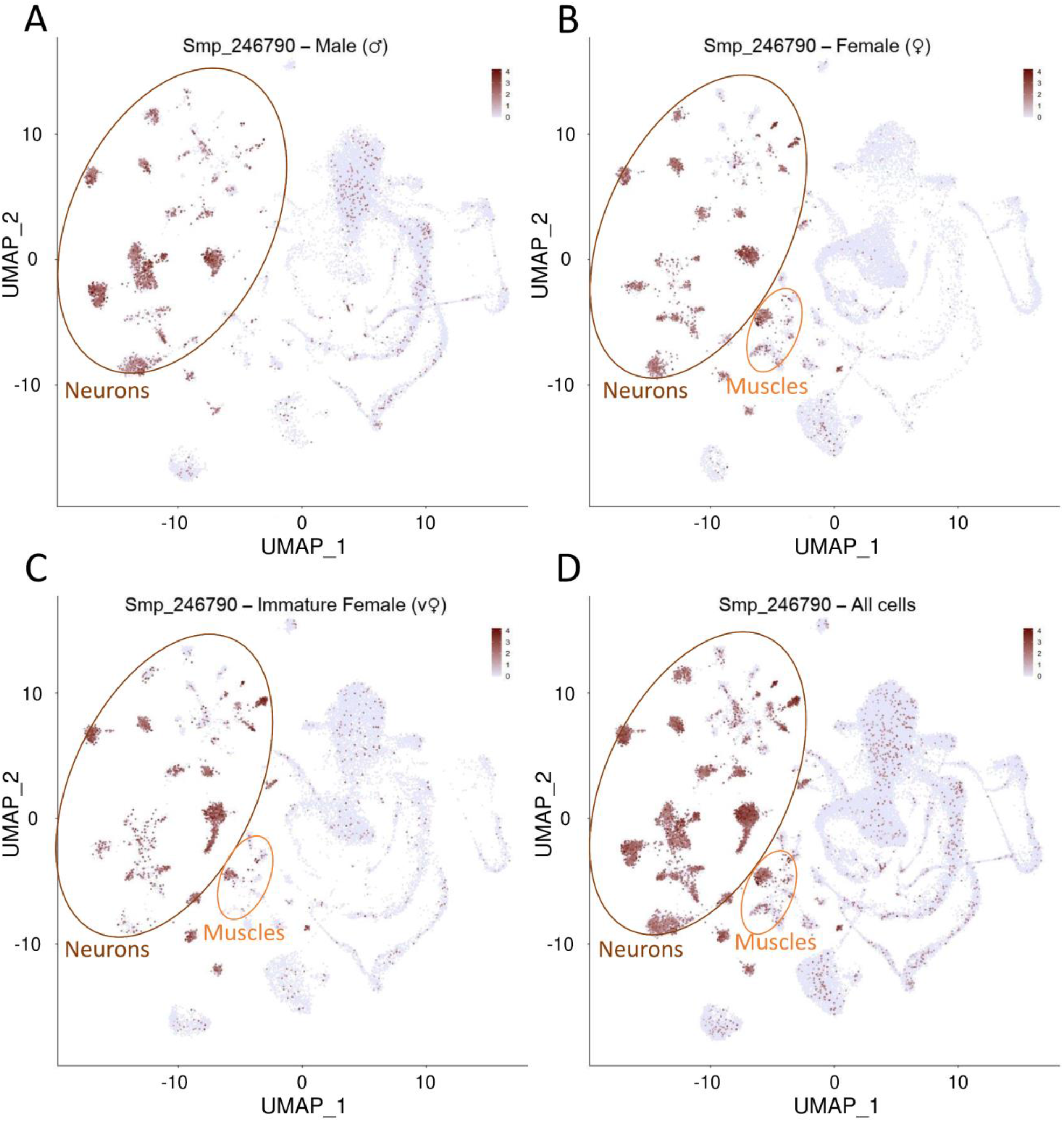
Cellular localization of *Sm.TRPM_PZQ_* expression in *S. mansoni.* **(A)** adult male, **(B)** adult female, **(C)** immature female, **(D)** overall sex and stages (SchistoCyte Atlas (*46*)). *Sm.TRPM_PZQ_* gene is essentially expressed in neurons for all sex and stages and is also expressed in muscle cells in females.

**Fig S6.**
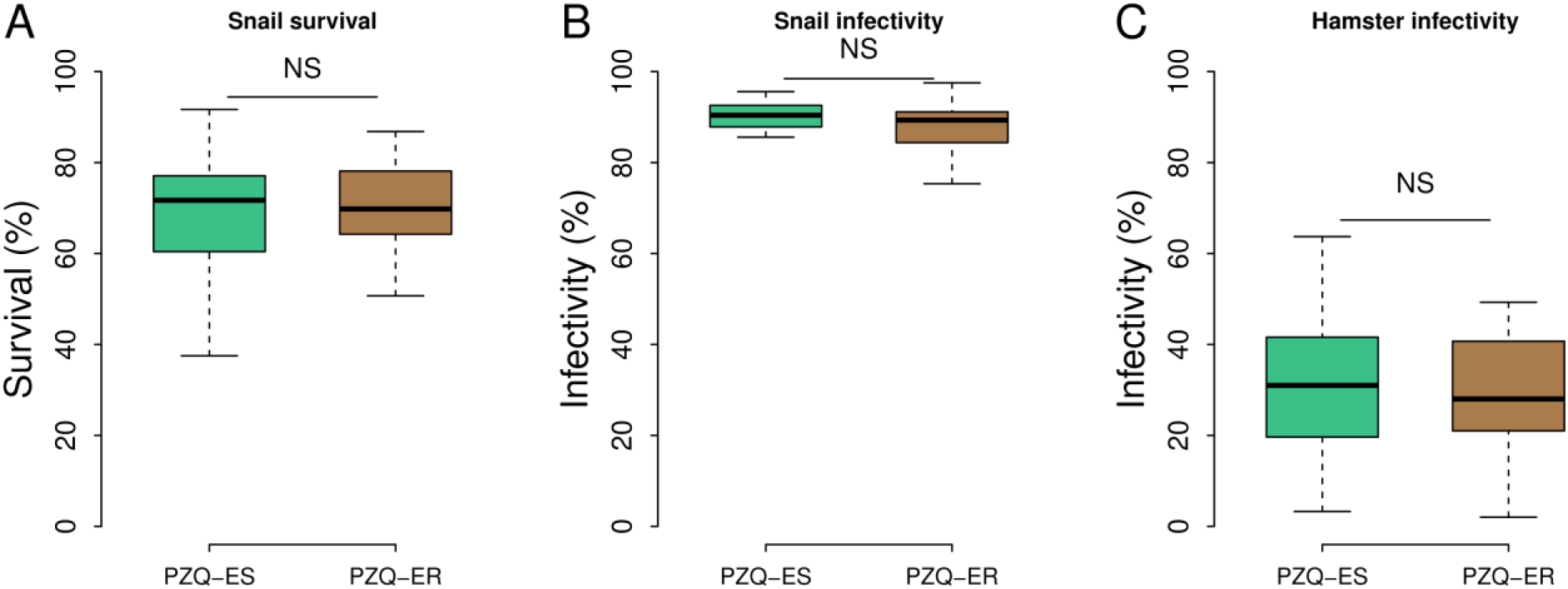
Fitness of SmLE-PZQ-ES and SmLE-PZQ-ER parasites. Comparison of several life history traits: **(A)** Snail survival (Welsh t-test, t = -0.662, p = 0.51), **(B)** infectivity to snails (Wilcoxon test, W = 123, p = 0.45), and **(C)** infectivity to hamsters (Welsh t-test, t = 0.725, p = 0.47) for 12 generations of SmLE-PZQ-ER and SmLE-PZQ-ES parasites. (NS: No significant difference between groups).

**Fig. S7.**
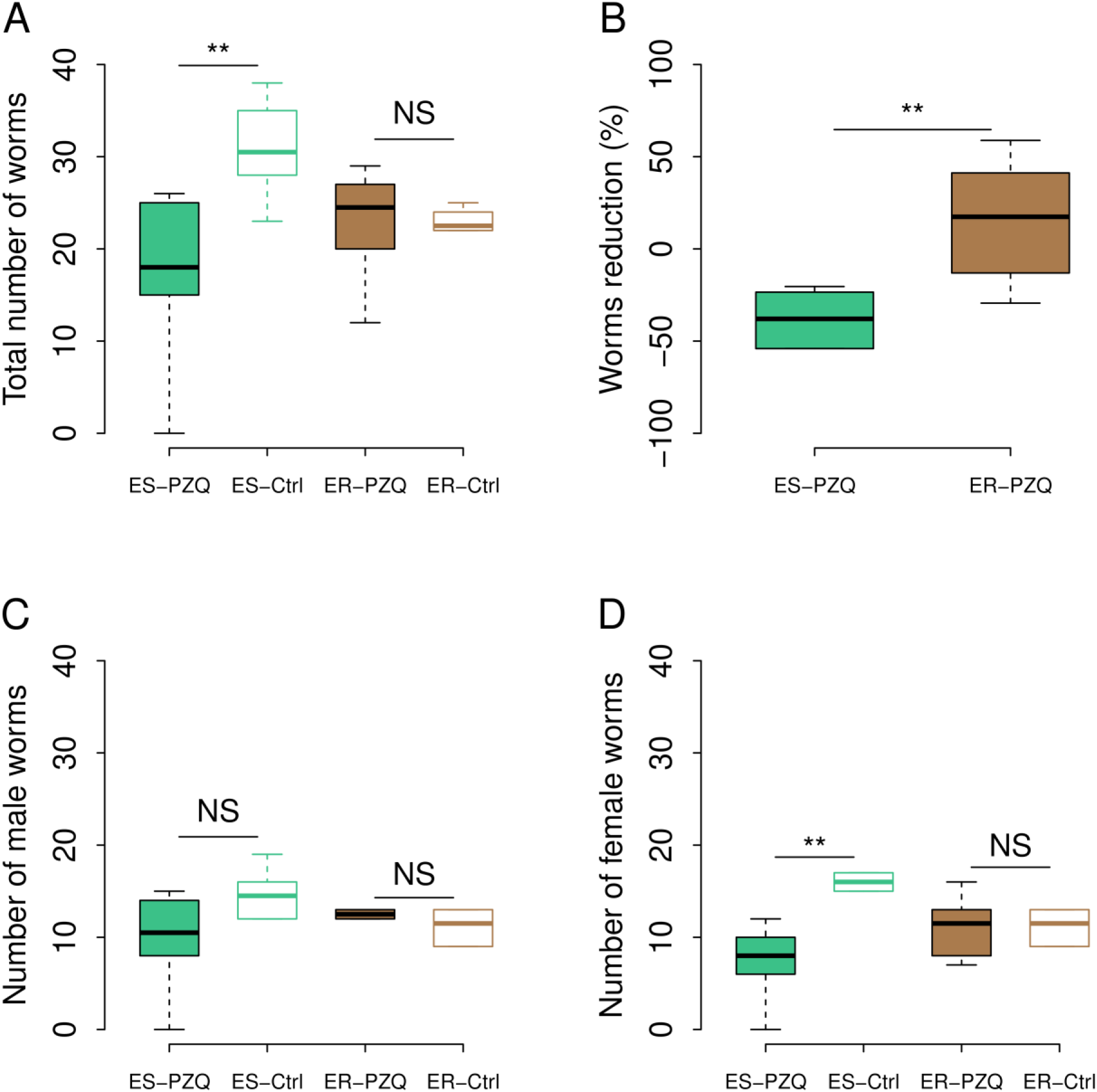
Impact of *in vivo* PZQ treatment on SmLE-PZQ-ER and SmLE-PZQ-ES parasites. Balb/c mice were infected with either SmLE-PZQ-ER or SmLE-PZQ-ES parasites populations and treated with 120 mg/kg of PZQ or DMSO (control group). **(A)** We observed no significant reduction in worm burden in SmLE-PZQ-ER parasites when comparing PZQ-treated and control (DMSO) treated animals (Wilcoxon test, p = 0.393). In contrast, we recovered significantly lower numbers of worms from PZQ-treated versus untreated mice infected with the SmLE-PZQ-ES parasite population (Wilcoxon test, p = 0.008). **(B)** The percent reduction observed was significantly different between the SmLE-PZQ-ES and SmLE-PZQ-ER parasites (Wilcoxon test, p = 0.0129). **(C)** While Interestingly, we did not reach significance for male worms (Wilcoxon test, p = 0.089), **(D)** we observed a large reduction in numbers of female worms recovered from PZQ-treated SmLE-PZQ-ES parasites relative to untreated animals (Wilcoxon test, p = 0.008) (*N* = 24 mice – 6 mice/group; NS: No significant difference between groups; \**p* < 0.05; ** *p* ≤ 0.02; *** *p* ≤ 0.002).

**Fig. S8.**
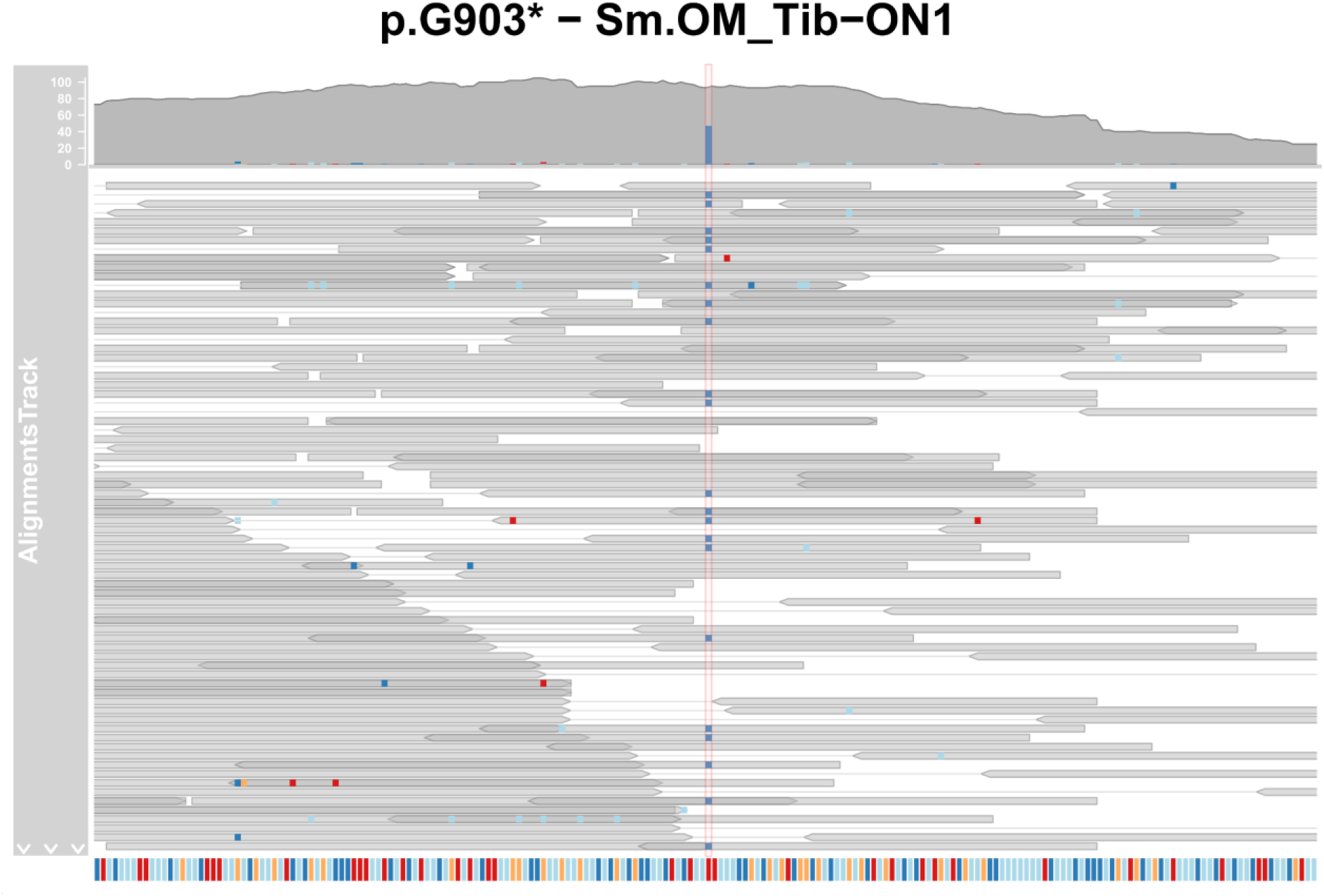
Stop codon identified in *S. mansoni* field sample from Oman. The stop codon p.G903* was identified in NGS exome capture data in one sample collected in Oman. This mutation occurred in exon 14 of the *Sm.TRPM_PZQ_* gene and the mutation is supported by a high number of reads.

**Fig. S9.**
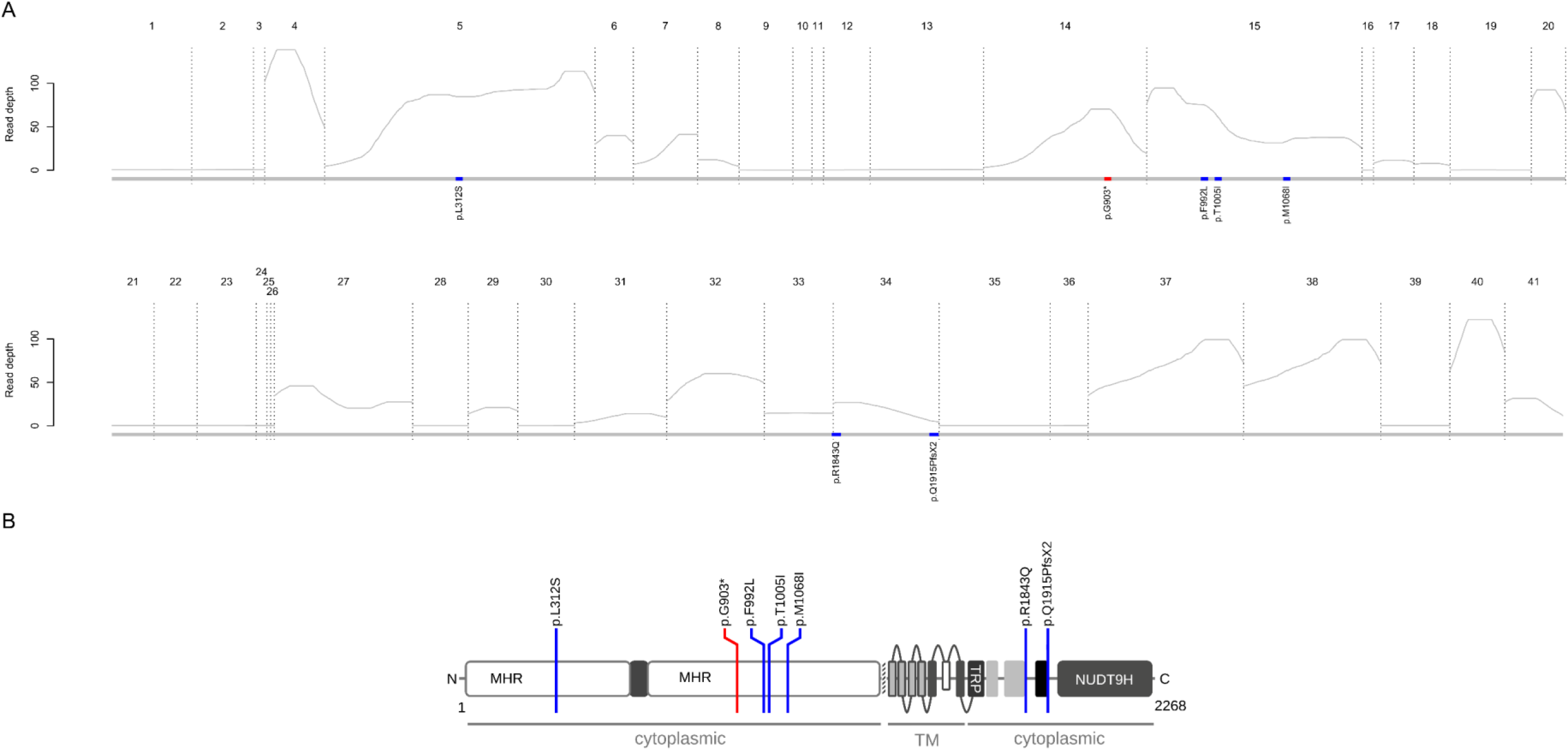
*Sm.TRPM_PZQ_* gene: average exon read depth and identified mutations in field samples. **(A)** Exons are numbered and delimited with dotted lines. Blue boxes on the grey line represent positions of the high frequency mutations. Red box represents the position of the low frequency resistant mutation. Mutation position mentioned below the boxes refers to the isoform 5. **(B)** Mutations were reported to the schematic structure of Sm.TRPM_PZQ_ modified from (*20*). The resistant mutation is located before the transmembrane domain (TM) leading to no a non-functional channel (MHR: TRPM homology region, TRP: TRP domain, NUDT9H: human ADP-ribose (ADPR) pyrophosphatase).

**Table S1. Genes in QTL regions on chr 2 and 3.**

*Separate file*

**Table S2.**
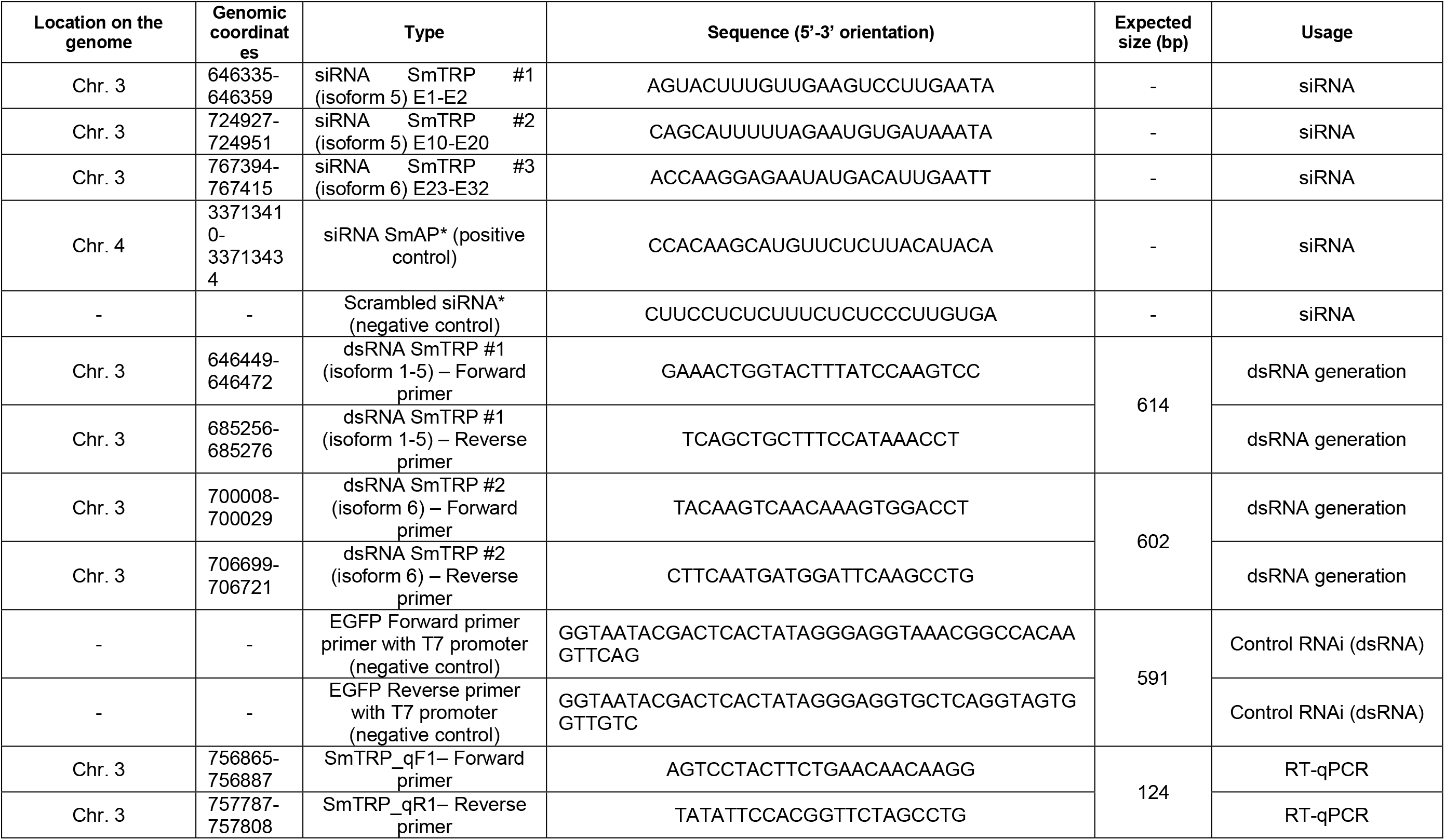
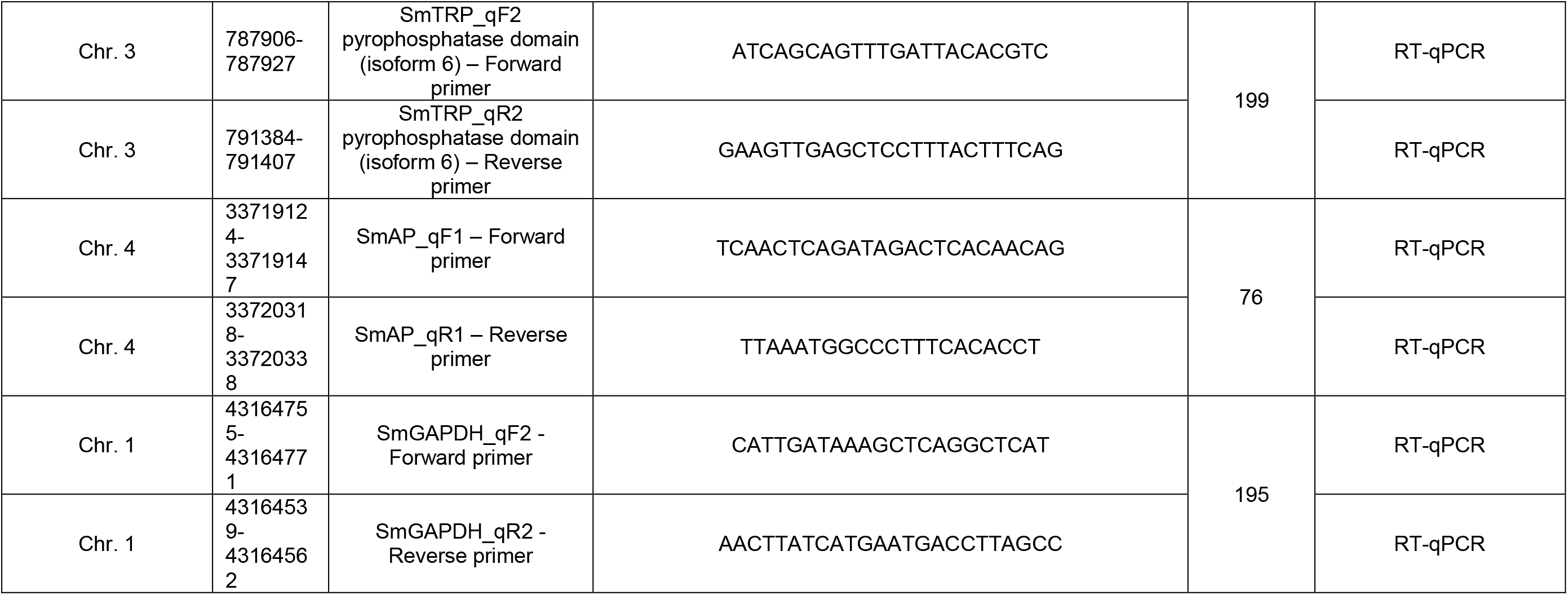
Summary table of RNAi for Sm.TRPM_PZQ._ siRNA sequences and primers used to generate dsRNA. Primer sequences used for RT-qPCR to quantify gene expression after RNAi treatment on worms (Chr.: Chromosome; E: Exon). * siRNA negative (scramble siRNA) and positive control (SmAP) have been used from Krautz-Peterson *et al.* (*70*).

**Table S3. Mutations present in *Sm.TRPM_PZQ_* in natural schistosome populations from 3 African countries (Senegal, Niger, Tanzania), the Middle East (Oman) and South America (Brazil).**

*Separate file*

**Table S4.**
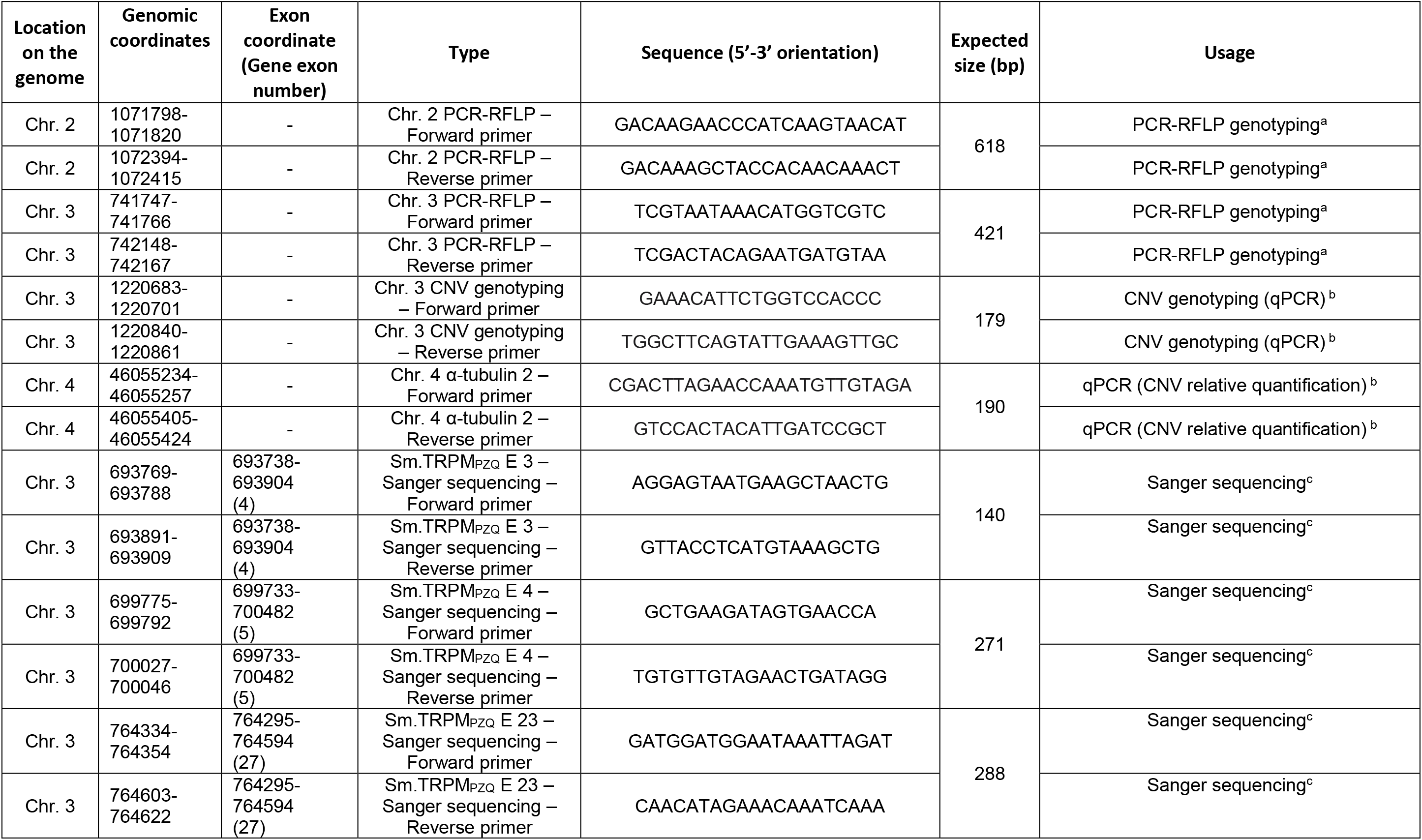
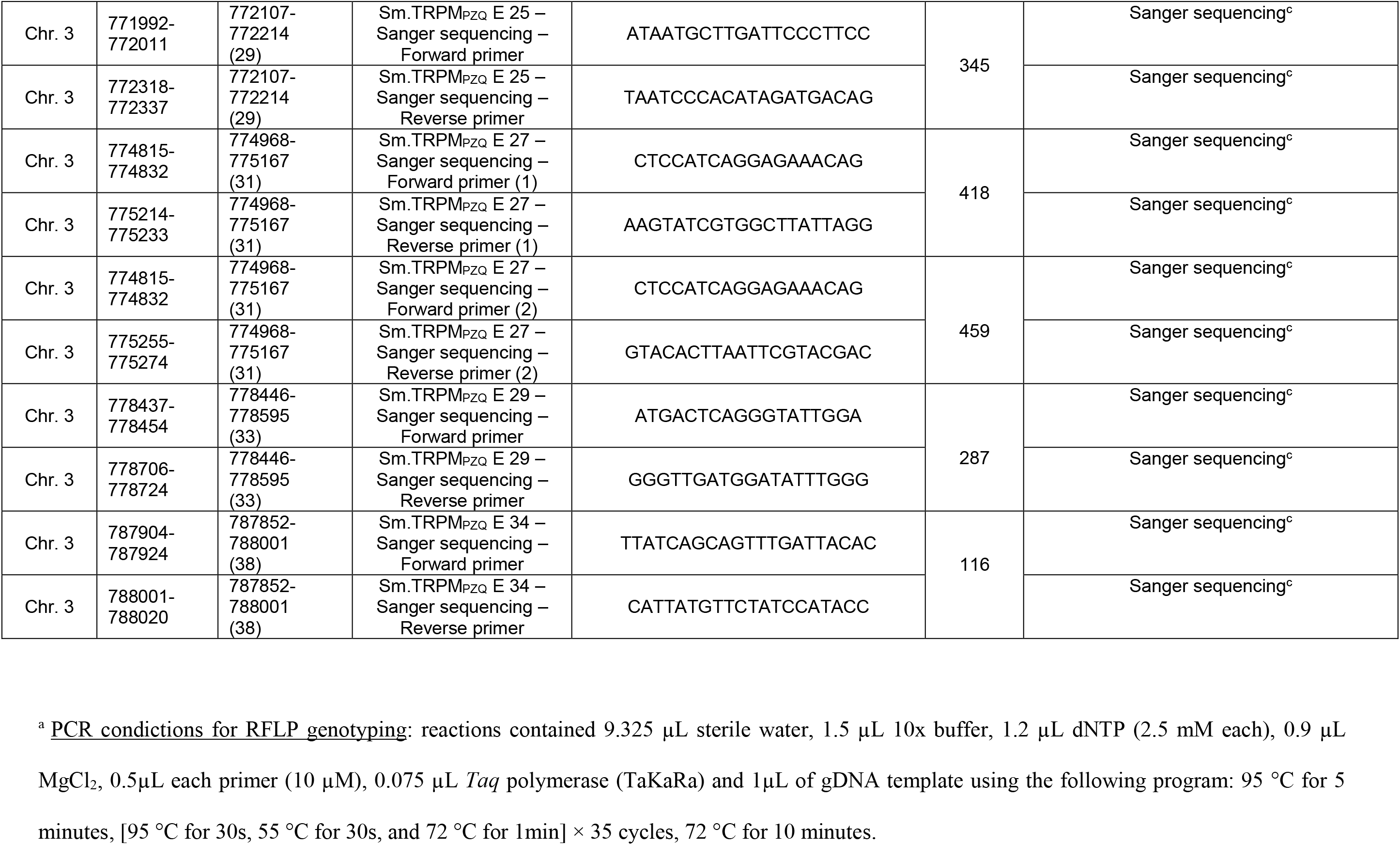
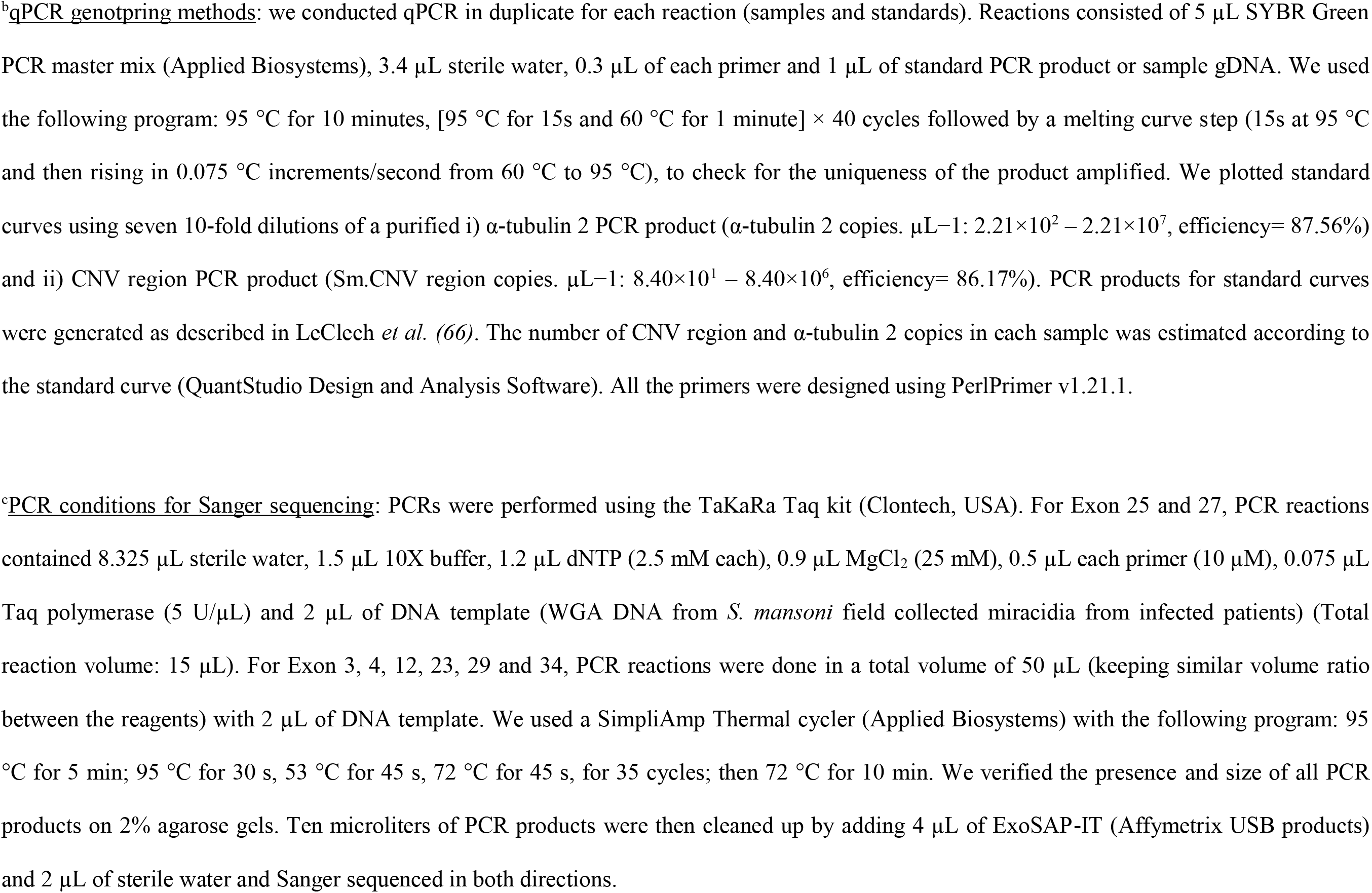
Summary table of all the primer sequences. used for i) PCR-RFLP and CNV quantification for single worm genotyping, ii) Sanger sequencing of Sm.TRPM_PZQ_ in field collected *S. mansoni* parasites (Chr. : Chromosome; E: Exon).

